# *CDKN1B* (p27^kip1^) enhances drug tolerant persister CTCs by restricting polyploidy following mitotic inhibitors

**DOI:** 10.1101/2024.02.20.581202

**Authors:** Elad Horwitz, Taronish D. Dubash, Annamaria Szabolcs, Ben S. Wittner, Johannes Kreuzer, Robert Morris, Aditya Bardia, Brian Chirn, Devon Wiley, Dante Che, Hunter C. Russel, Xcanda Ixchel Herrera Lopez, Douglas B. Fox, Ezgi Antmen, David T. Ting, Wilhelm Haas, Moshe Sade-Feldman, Shyamala Maheswaran, Daniel A. Haber

## Abstract

The mitotic inhibitor docetaxel (DTX) is often used to treat endocrine-refractory metastatic breast cancer, but initial responses are mitigated as patients eventually have disease progression. Using a cohort of *ex vivo* cultures of circulating tumor cells (CTCs) from patients with heavily pretreated breast cancer (n=18), we find two distinct patterns of DTX susceptibility, independent of clinical treatment history. In CTCs cultured from some patients, treatment with a single dose of DTX results in complete cell killing, associated with accumulation of non-viable polyploid (≥8N) cells arising from endomitosis. In others, a transient viable drug-tolerant persister (DTP) population emerges, ultimately enabling renewed proliferation of CTCs with preserved parental cell ploidy and DTX sensitivity. In these CTC cultures, efficient cell cycle exit generates a ≤4N drug-tolerant state dependent on *CDKN1B* (p27^Kip1^). Exposure to DTX triggers stabilization of CDKN1B through AKT-mediated phosphorylation at serine 10. Suppression of *CDKN1B* reduces the number of persister CTCs, increases ≥8N mitotic cells and abrogates regrowth after DTX exposure. Thus, CDKN1B-mediated suppression of endomitosis contributes to a reversible persister state following mitotic inhibitors in patient-derived treatment refractory breast cancer cells.

**Summary in bullets:**

- Transient DTX tolerant persister cells emerge in some patient-derived cultured CTCs.
- DTX-tolerant persisters restrict endoreduplication and polyploidy through CDKN1 (p27^kip^^1^).
- DTX exposure induces CDKN1B stabilization through AKT mediated phosphorylation at serine 10.
- Suppression of polyploidy underlies a drug tolerant persister state specific to mitotic inhibitors.

## Introduction

Cancer-associated genomic instability is thought to underly the rapid emergence of mutations, leading to acquired resistance to both cytotoxic and targeted therapies. However, as cancers evolve, cellular plasticity, and additional non-genetic factors contribute to clinical treatment failure. Initially proposed in the context of targeted therapy for *Epidermal Growth Factor Receptor (EGFR)*-mutant non-small cell lung cancer^1^, the emergence of Drug Tolerant Persisters (DTP) has been extended to other therapeutic agents^2–5^. Instead of comprising a pre-existing population of mutants selected by drug exposure^6, 7^, these cells appear to be induced by cellular stressors, producing a transient population of slowly proliferating, drug-resistant cells. Under continuous drug selection pressure, DTPs may acquire a genetic mutation mediating permanent drug resistance, or alternatively they may revert to a proliferative drug-sensitive state in the absence of selection^1^. The fundamental mechanisms responsible for the appearance and maintenance of DTPs remains uncertain, with most studies focused on markers of cell states within established DTPs, independent of specific drug-induced signals that trigger this phenomenon. These include alterations in chromatin state rendering DTPs sensitive to inhibitors of histone deacetylases and Insulin Growth Factor Receptor^1, 8^(IGFR); disruption of fatty acid oxidation leading to susceptibility to GPX4 inhibition^4, 9^; sensitivity to suppressors of YAP-TAZ^10^, Fibroblast Growth Factor Receptor^11^ (FGFR) and CDK9-dependent transcriptional regulation^2^. Correlations have also been proposed between established DTPs and autophagy modulation^12^ and a developmental cell state called diapause^2^. While these observations point to a cell state that may share characteristics of quiescence, the acute drug-induced signals that trigger such a state are not understood.

The taxane class of mitotic inhibitors are widely applied as anti-tumor agents in breast, lung and prostate cancers, where they are often used in patients with advanced disease that has acquired resistance to first line therapeutic agents. Taxanes bind to microtubules and inhibit their depolymerization, preventing mitotic progression from metaphase to anaphase and triggering cell death^13^. As a class, taxanes target rapidly dividing cells, but cancers eventually resume proliferation upon acquiring drug resistance. Reported mechanisms of resistance, derived primarily from model systems, include activation of drug metabolizing pathways or drug efflux pumps, mutations targeting drug binding domains within tubulin proteins, increased expression of distinct β-tubulin subunits and upregulation of apoptosis inhibitors, including Mcl1, Bcl2 and BclXl^14–19^. However, the relative contribution of these drug resistant mechanisms to clinical treatment failure is uncertain, particularly since taxanes are often administered to patients with advanced, heavily pre-treated cancers, whose apparent cell plasticity is not well recapitulated in most model systems. Clinical reports of successful retreatment with taxanes in cancers that had previously progressed following earlier therapeutic courses^20^ suggest possible contributions of reversible drug resistance phenotypes, yet these are not well defined.

Circulating tumor cells (CTCs) are shed from tumor deposits into the bloodstream and thus provide an opportunity to non-invasively sample advanced cancer cells in the context of their evolving sensitivity and resistance to therapeutic interventions^21–26^. While many CTCs in the bloodstream undergo cell death, a subset are viable metastatic precursors, which may be propagated *in vitro* following gentle microfluidic cell enrichment technologies^22, 27^. We have established a cohort of such patient CTC-derived cultures from blood specimens of women with advanced, heavily treated hormone receptor positive (HR+) and triple negative (TNBC) breast cancers. These long-term *ex vivo* cultures of CTCs, which are maintained under anchorage independent, hypoxic and stem-cell like culture conditions, are tumorigenic in immune deficient mice and they display features of cellular plasticity, consistent with advanced cancer cells^22, 28, 29^. Here, we use cultured CTCs from a cohort of 18 patients with treatment-refractory metastatic breast cancer, modeling their sensitivity to DTX. We describe heterogeneity across different patient-derived cultures, with some being highly susceptible to DTX, while others undergo a reversible DTP-like state, dependent upon drug-induced stabilization of the cyclin dependent kinase (CDK) inhibitor CDKN1B.

## Results

### Heterogeneity in the generation of DTX-tolerant persister cells in patient-derived breast CTC cultures

We tested 18 different breast cancer CTC cultures, each generated from a patient with advanced metastatic-breast cancer who had received multiple serial courses of both standard of care and experimental therapies (Figure 1A, Supplementary table 1). Fourteen CTC-cultures were derived from patients whose breast cancer was HR+ at presentation and three from patients with TNBC (Supplementary Table 2). Among all CTC cultures, 11 were derived from patients who had previously been treated with DTX or other taxane-based mitotic inhibitors such as Docetaxel, Paclitaxel, Navelbine, Nab-paclitaxel and Eribulin as part of their clinical care (median duration of response 3.5±2.79, SD, Figure 1A and Supplementary Table 1). Six patients had two or three iterations of treatment courses using mitotic inhibitors (Figure 1A and Supplementary table 1). Of note, the duration of treatment with each iteration of taxane drug therapy, which is a measure of clinical responsiveness, did not significantly decrease in second or third line treatment courses (Supplementary Figure 1A). This serial intermittent sensitivity to taxanes raises the possibility that treatment failures may not have resulted from stably acquired mutational mechanisms.

**Figure 1:**
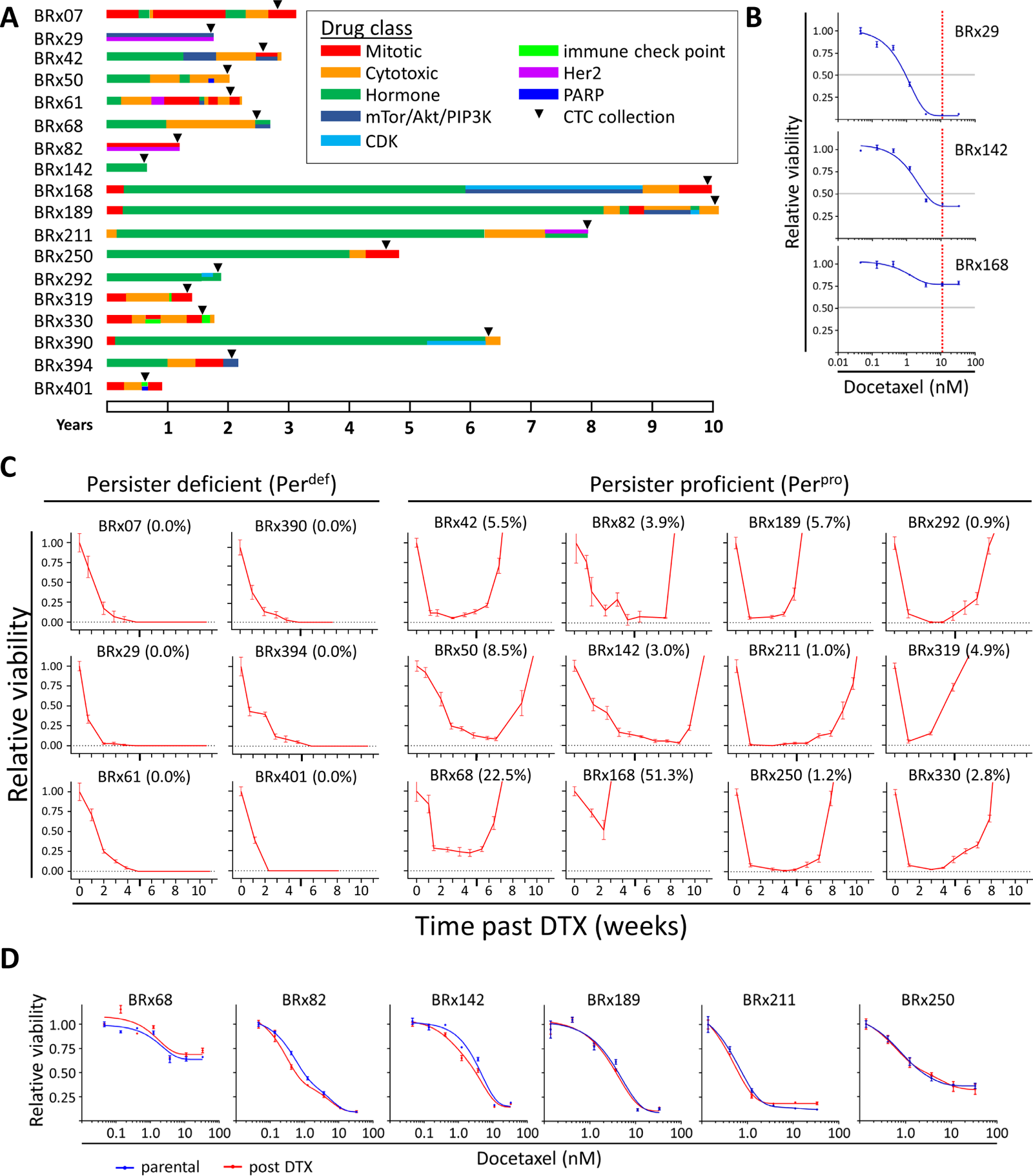
Heterogeneity in the generation of DTX-tolerant persister cells in patient-derived breast CTC cultures. **1A:** Treatment histories of 18 patients with metastatic breast cancer from whom CTCs were cultured *ex vivo*. BRx-319, BRx-330 and BRx-401 were derived from patients with Triple Negative Breast Cancer (TNBC), while all other cultures were derived from patients with Hormone Receptor positive (HR+) breast cancer. Treatment histories for some patient-derived CTC lines have been previously described: BRx-07, BRx-42, BRx-50, BRx-61 and BRx-68^22^, BRx-29^77^, BRx-82 and BRx-142^29^, BRx-211, BRx-250 and BRx-394^78^. **1B:** Representative CTC lines (BRx-29, BRx-142 and BRx-168) treated with increasing concentrations of DTX for 6 days show differences in residual viable cell populations (red line). Cell viability at each dose was measured in biological triplicates. **1C:** Regrowth of CTCs following pulse DTX (10nM, 16 hours) in per^pos^ cultures (N=12) but not others per^def^ cultures (N=6) within 10 weeks after drug exposure. A representative (from 2-3 biological replicates) longitudinal cell viability curve based on cell count is shown for each breast CTC culture, with each point representing the mean ± SEM (n≥3 experimental replicates). BRx refers to the name of all CTC lines. Parentheses denote the percent of remaining viable cells at the lowest point in longitudinal monitoring. **1D:** Restored parental cell DTX sensitivity in cultured CTCs emerging from DTX-tolerant persister populations. Six representative per^pro^ CTC cultures were expanded following their regrowth after pulse DTX (10nM, 16 hrs), tested with increasing doses of DTX (red), and compared to matched untreated parental cells (blue). Cell viability at each dose was measured across biological triplicates.

Breast CTC cultures isolated from different patients exhibit highly variable responses to increasing doses of DTX, with some cultures displaying complete cell killing at higher drug doses, while others reach a plateau with minimal residual fractions (1-E_max_) ranging from 6% to 80% at DTX concentrations of up to 30nM for 6 days (Figure 1B and Supplementary Figure 1B). The distinct patient-specific response patterns to DTX by CTC cultures are highly reproducible and independent of the length of time they were cultured *in vitro* since isolation from blood specimens. To further model these patterns and allow viable isolation of drug-tolerant persisters (DTP), we first applied a single 16-hour pulse of 10nM DTX, a dose that achieves maximal cell killing across all CTC cultures after 6 days (Supplementary Figure 1B, 1C) washed the cells and let them grow in a drug-free media for 11 weeks. Six CTC cultures are completely eradicated under these conditions, with no viable cells counted at a median time of 5 weeks past DTX pulse (range: 2-6 weeks) and no rare colonies re-emerging for up to 11 weeks (Figure 1C). However, in the other 12 CTC cultures, a fraction of cells remain non-proliferative yet viable (median: 4.4% surviving cells, range: 0.9-51.3%; Figure 1C) for an extended period (median 7 weeks after DTX pulse; range 3-9 weeks) after which they resume proliferation (Figure 1C). We call CTC cultures capable of producing such drug-tolerant persisters “persister-proficient” (per^pro^), and those unable to produce such cells “persister-deficient” (per^def^).

The ability of CTCs to regrow after a pulse of 10nM DTX is correlated with other parameters of drug sensitivity when cells where cultured with different DTX concentrations for 6 days (Figure 1B and Supplementary Figure 1B). Per^def^ BRx cultures exhibit lower area under the curve (AUC) values (median: 0.0088, range: 0.0031-0.0131; AUC= relative viability × nM DTX) compared to AUC values in per^pro^ cultures (median: 0.0148, range: 0.0065-0.0261, p<0.05; Supplementary Figure 1D, upper panel). Differences in IC_50_ values were not readily calculated, since 4 of 12 per^pro^ CTC cultures did not achieve 50% cell death following DTX (Supplementary Figure 1D, middle panel). However, the per^def^ CTC cultures have lower fractions of surviving cells 7 days after DTX treatment (median 1-E_max_ = 14.5% ± 0.055%, range 5.8 to 37.4%) compared with per^pro^ CTC cultures (median 1-E_max_ = 37.7%±19.3%, range: 10.7-76.9%, p<0.05, Supplementary Figure 1D, lower panel).

Clinical treatment histories of patients from whom the CTC cultures had been generated do not reveal a correlation between previous taxane exposure and susceptibility to DTX *in vitro*. Five of the six patients whose CTCs show high sensitivity to DTX (per^def^) had received taxanes as part of their care prior to CTC isolation and were hence considered to have acquired drug resistance in the clinical setting. Of the 12 patients whose CTC cultures showed regrowth (per^pro^) after pulse DTX, eight had received taxane therapy prior to CTC isolation, while the other four patients had not been treated with mitotic inhibitors (Figure 1A and Supplementary Table 1). The susceptibility of cultured patient-derived CTCs to DTX-mediated cell killing is thus not a strict reflection of clinical treatment history, suggesting intrinsic differences in drug responses across the distinct parental tumors. These differences extend to both HR+ breast cancer and TNBC histologies.

Remarkably, the DTX sensitivity profiles of CTC cultures that ultimately resume proliferation after receiving pulse DTX are identical to those of their respective parental cells (Figure 1D). These cells thus appear to have undergone a transient DTP state rather than acquiring stable, heritable drug resistance.

### Proliferative arrest and ploidy changes distinguish per^def^ from per^pro^ CTCs

To investigate early drug-induced changes that may precede the generation of DTPs, we first compared expression profiles of CTCs under baseline culture conditions versus CTCs that remained viable four days after pulse DTX. We performed RNA sequencing at the single cell level to address potential heterogeneity in CTC cultures, comparing two per^pro^ cultures (BRx-82 (94 cells) and BRx-142 (36 cells)) with two per^def^ cultures (BRx-29 (94 cells) and BRx-394 (44 cells)). Remarkably, UMAP analysis shows distinct drug treatment-associated patterns between per^pro^ and per^def^ cultures (Figure 2A, upper panel). DTX-treated per^pro^ cultures exhibit a synchronous separation between treated and non-treated cells, as demonstrated using nearest neighbor analysis (BRx-82: 84.7% treated or untreated cells are near their respective cell type, p<10^-5^; BRx-142: 86.1%, p<0.001; Figure 2A, lower panel). In contrast, DTX-treated and untreated per^def^ CTC cultures remain randomly mixed and fail to show any clear drug-induced separation (BRx-29: 59.8%, p=0.076; BRx-394: 40.9%, p=0.87; Figure 2A, lower panel). This suggests a more uniform transcriptional response to DTX in per^pro^, compared with per^neg^ CTCs.

**Figure 2:**
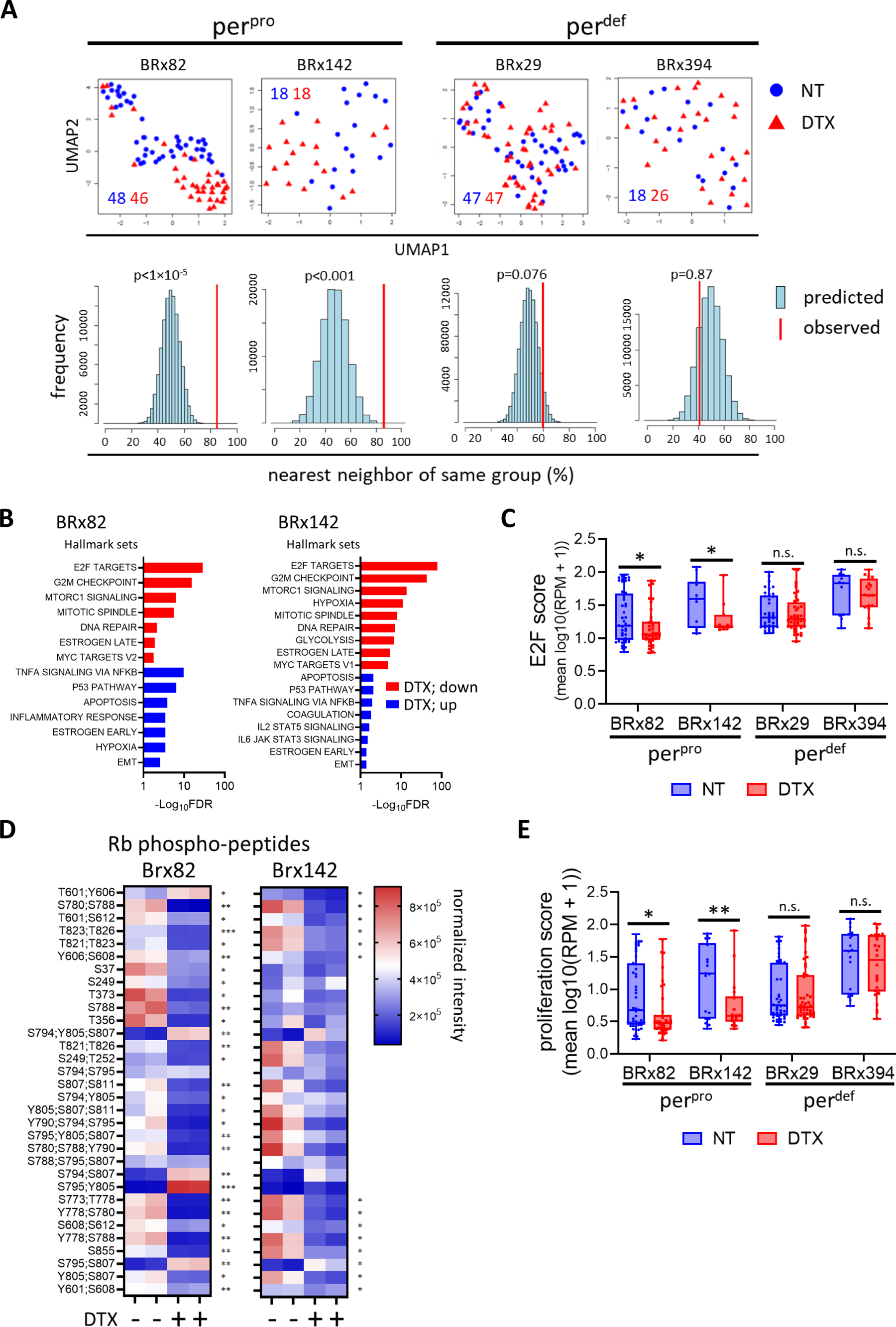
Proliferative arrest and ploidy changes distinguish per^def^ from per^pro^ CTCs. **2A:** UMAPs and nearest neighbor analysis of of single CTCs from two per^pro^ (BRx-82 and BRx-142) CTC cultures and two per^def^ CTC cultures (BRx-29 and BRx-394) at day four following DTX (10nM) exposure. ***Upper panel***: UMAP pattern of single cell RNA-seq from untreated (blue circles) and DTX-treated (red triangles) CTCs, showing segregation pattern in per^pos^, but not per^def^ cultures. The number of cells sequenced for each condition is shown and color coded. ***Lower panel***: computational analysis (nearest neighbor; see methods) for each cell line shown in UMAP using permutation testing, showing that a large fraction of non-treated *versus* DTX-treated per^pro^ CTCs segregate according to their respective grouping and outside of the random prediction distribution (BRx-82: 84.7%, p=10^-5^; BRx-142: 86.1%, p<0.001), in contrast to per^def^ CTC cultures, where grouping treated *versus* untreated cells overlaps with a predicted random distribution (BRx-29: 59.8%, p=0.076; BRx-394: 40.9% p=0.87). The red bar denotes the observed percent of cells whose nearest neighbor is from the same grouping (treated *versus* untreated); the predicted random distribution (blue) is derived from a hundred thousand random iterations of data from each CTC culture. **2B:** GSEA pathway analysis of genes differentially expressed in single cell RNA-seq of untreated and DTX-treated per^pro^ CTC cultures, showing suppression of proliferation-associated pathways in DTX-treated CTCs. **2C:** E2F signaling metagene score, demonstrating suppression in DTX-treated per^pro^ CTCs (BRx-82, BRx-142), but not in per^def^ CTC culture (BRx-29, BRx-394). P-value calculated using two-tailed Student’s T-test (* p<0.05). The individual values of each cell are overlaid on the boxplot. **2D:** Multiplexed quantitative mass spectrometric-based phosphoprotemics analysis of untreated and DTX-treated per^pro^ CTCs (BRx-82, BRx-142), showing decline in multiple RB1 phosphopeptides. P-value calculated using two-tailed Student’s T-test (* p<0.05, ** p<0.01, *** p<0.001). **2E:** Proliferative signature^79^ in DTX-treated per^pro^ CTCs, and per^def^ CTC cultures. P-value calculated using two-tailed Student’s T-test (* p<0.05, ** p<0.01). The individual values of each cell are overlaid on the boxplot.

GSEA analysis of genes differentially expressed between untreated and DTX-treated per^pro^ cells (BRx-82 and BRx-142) shows strong suppression of cell proliferation across both CTC lines (E2F targets, G2/M checkpoint genes, mTORC1 signaling; BRx-82 FDRs from 4.07x10^-29^ to 4.76x10^-7^; Brx-142 FDRs from 9.03x10^-79^ to 2.64x10^-14^; Figure 2B). In contrast, analysis of untreated and DTX-treated per^def^ cultures (BRx-29, BRx-394), does not reveal any commonly shared induced or suppressed pathways (Supplementary Figure 2A). An E2F metagene signature (derived using E2F target genes collected from the Hallmark E2F target gene set) assigned to each cell within untreated and DTX-treated cultures also shows significant suppression of the E2F pathway in per^pro^ cultures but not in per^def^ cells (p<0.05; Figure 2C).

Multiplexed quantitative phosphoproteomic mapping^30^ of untreated and DTX-treated per^pro^ CTC cultures (BRx-82 and BRx-142) shows significant (p<0.05) decrease in RB1 RB1 (a key regulator of activator E2F function^31^) phospho-peptides representing 23 unique phospho-sites (30/32 peptides in BRx-82; 14/32 peptides in BRx-142; Figure 2D). Coincident with the robust decline of E2F targets and Rb phosphorylation, DTX-treatment significantly suppresses proliferation signatures in per^pro^ CTCs but not in per^def^ CTCs (p<0.05 for BRx-82, p<0.01 for BRx-142, Figure 2E). Together, these findings show that per^pro^ and per^def^ CTC cultures exhibit differential proliferative responses to DTX exposure, with a robust decline of the E2F-RB1 proliferative signaling axis in the per^pro^ cells.

### Increased endomitosis following DTX in per^neg^ CTCs

To determine the consequences of the differential cell cycle arrest between per^pro^ and per^def^ CTC cultures, we applied flow cytometry using the mitotic marker phospho-Histone H3 (Serine 10, pHH3) to assess M phase and Hoechst 33342 DNA dye to quantitate ploidy (Methods). In per^pro^ (n=12) and per^def^ (n=6) CTC cultures, we measured the fraction of mitotic cells as a function of DNA ploidy, both at untreated baseline and 7 days after DTX exposure. As expected, DTX treatment results in a shift from 2N DNA content to 4N and 8N (Figure 3A). The total number of cells at these different ploidy levels is comparable between per^pro^ and per^def^ CTC cultures at baseline. Remarkably, however, per^def^ CTCs exhibit considerably more mitotic ≥8N cells (2.88 ± 0.66% pHH3+ of all live cells) compared with per^pro^ CTCs after DTX treatment (0.79 ± 0.22%, p<0.001; Figure 3B). In contrast to these polyploid mitotic cells, CTCs with 4N ploidy following DTX showed no significant differences in mitotic activity between per^pro^ and per^def^ cultures (Supplementary Figure 3A). Further, ploidy assessment of DTX-treated persister cells resuming proliferation (per^pro^) shows a similar profile to their respective untreated parental cells, with a preponderance of 2N ploidy (Figure 3C). Taken together, these studies indicate that per^def^ CTCs undergo an incomplete cell cycle arrest following drug exposure, reaching a ploidy of ≥8N and continuing to enter mitosis with ultimate loss of cell viability. The more synchronized and complete cell cycle arrest in per^pro^ CTCs results in a viable persister population, which subsequently re-enters the cell cycle with ploidy and DTX drug sensitivity similar to that of parental populations.

**Figure 3:**
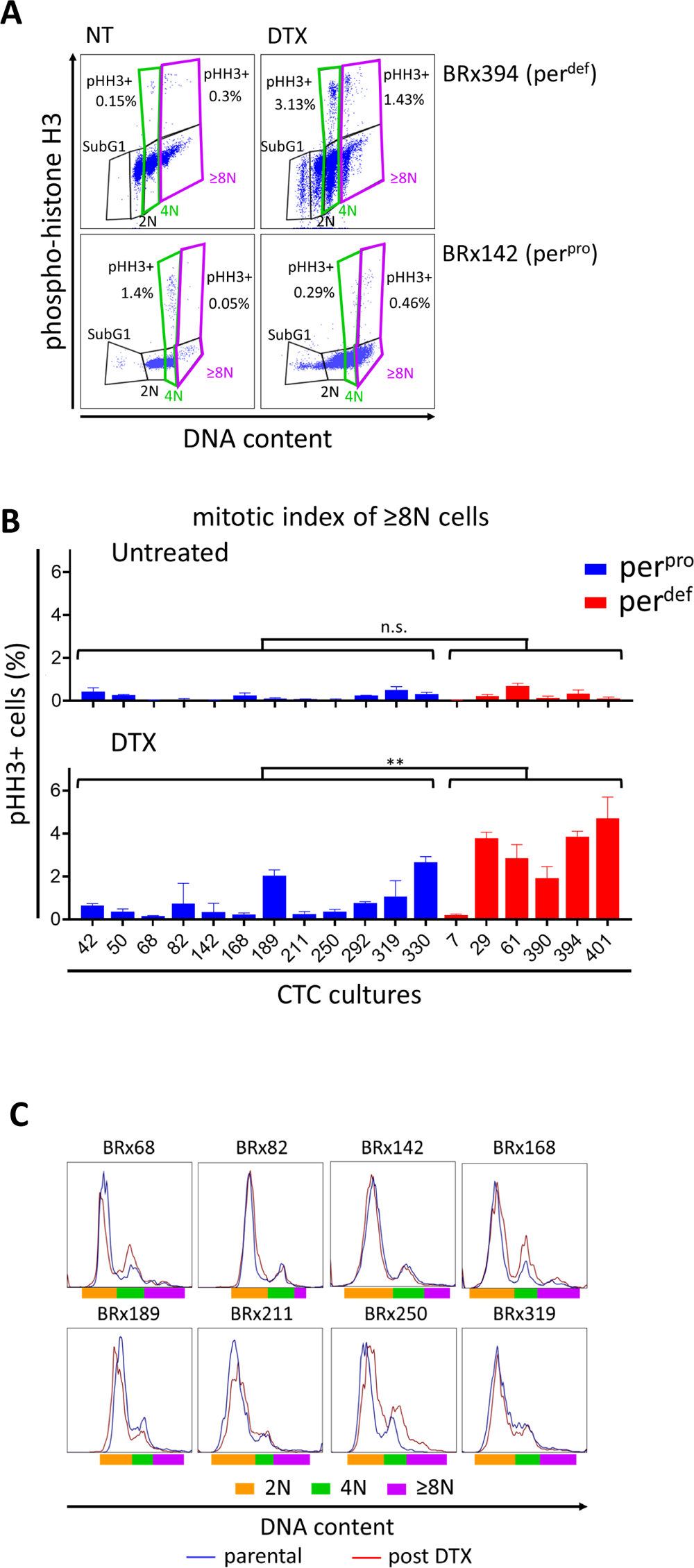
Increased endomitosis following DTX in per^neg^ CTCs. **3A:** Representative scatter plot (flow cytometry), demonstrating the fraction of per^def^ (BRx-394) and per^pos^ (BRx-142) CTCs with mitotic activity (phospho-histone H3; pHH3) as a function of DNA content (2N: black, 4N: green, >8N: purple) at day 7 following pulse DTX (10 nM) versus untreated controls (NT). **3B:** Quantification of pHH3-positive CTCs with ≥8N ploidy, at untreated baseline or at day 7 following puse DTX. Multiple per^pro^ (blue) CTC cultures and per^def^ (red) CTC cultures are shown, with accumulation of ≥8N mitotic cells in per^def^ cultures. CTC cultures are listed by their individual BRx number). Bar graphs represent mean ± SEM of three biological repeats, with p-value was calculated using two-tailed Student’s T-test (** p<0.01). **3C:** DNA content analysis (flow cytometry, Hoechst 33342 staining) of CTC cultures that re-established proliferation from per^pro^ DTPs three months after pulse DTX (10 nM), showing overlapping ploidy between the recovered treated CTCCs (red) and their matched untreated parental cells (blue). Ploidy is denoted as 2N: orange, 4N: green, ≥8N: purple. Shown is one of two biological repeats.

### DTX-induced stabilization of CDKN1B restricts endoreduplication and enhances DTPs

Since DTX-induced proliferation arrest distinguishes between per^pro^ and per^def^ CTC cultures, we tested whether differential regulation of CDK inhibitors contributes to this phenotype using bulk RNA sequencing. At baseline, per^pro^ CTC lines (n=12) and per^def^ CTC lines (n=6) display no appreciable differences in the expression of *CDKN1A* (p21) and *CDKN1B* (p27), both of which are present at significant levels (*CDKN1A*: median 150 ± 50 RPM, *CDKN1B*: 53 ± 11 RPM), whereas expression of *CDKN1C*, *CDKN2A*, *CDKN2B*, *CDKN2C*, and *CKDN2D* is negligible (<10 RPM, Supplementary Figure 4A). Although *CDKN1A* (p21) mRNA is highly expressed in the CTC cultures, siRNA mediated knockdown of CDKN1A in per^pro^ CTCs (BRx-82) does not affect their response to DTX treatment (Supplementary Figures 4B and 4C). In contrast, *CDKN1B (p27)* expression is induced in per^pro^ CTCs (BRx-142, BRx-82) following DTX exposure, as determined by both mass spectrometry and western blot analysis (Supplementary Figures 4D and E). At the single cell level, flow cytometry identifies an increased subpopulation of *CDKN1B-*overexpressing cells (CDKN1B^high^) within per^pro^ CTC cultures (BRx-68, BRx-82, BRx-142, BRx-211, BRx-250, BRx-292) 7 days after DTX exposure (median fold increase = 2.4 ± 0.79, range 1.7 to 7.0; Figures 4A and 4B). DTX-treated per^def^ CTCs do not yield sufficient viable cells to quantify rare CDKN1B^high^ subpopulations.

**Figure 4:**
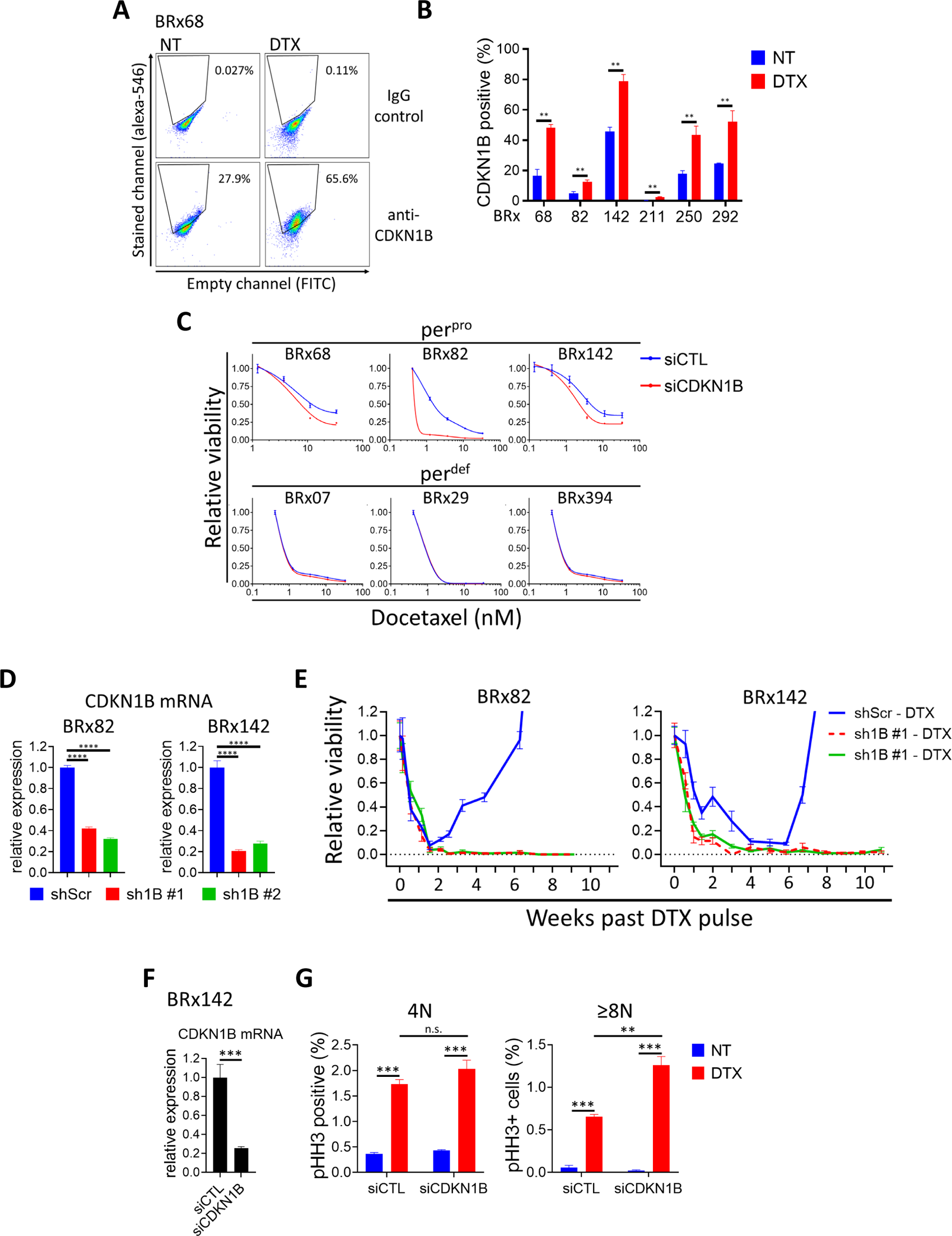
DTX-induced stabilization of CDKN1B restricts endoreduplication and enhances DTPs. **4A:** DTX treatment upregulates CDKN1B expression in a fraction of CTCs. Representative scatter plot showing untreated (NT) and DTX-treated (10nM, day 7) CTCs, stained with antibody against CDKN1B and analyzed by flow cytometry. **4B:** Bar graph: (mean ± SEM) increase in CDKN1B-positive cells in DTX-treated per^pro^ CTCs (BRx-68, BRx-82, BRx-142, BRx-211, BRx-250, BRx-292), compared with untreated cells (NT). n=3 biological repeats, with p-value calculated using two-tailed Student’s T-test (** p<0.01). **4C:** Suppression of DTPs in per^pro^ CTCs following *CDKN1B* knockdown using transient siRNA transfection. CTC cultures were transfected with siCDKN1B (red) or siControl (blue), and after three days treated with increasing concentrations of DTX (Knockdown efficiency is show in Supplemetary Figure S4F). Knockdown of *CDKN1B* decreases the minimal residual fraction of viable CTCs in per^pro^ CTC cultures (BRx-68, BRx-82, BRx-142), but it has no effect in the already low viable cell fraction in per^def^ CTC cultures (BRx07, BRx-29, BRx-394). 1-E_max_ at each DTX dose was measured (n= 3 biological repeats). p-value (p<0.0001 for each of the BRx-68, BRx-82 and BRx-142 CTCs) was calculated using a two-tailed Student’s T-Test. **4D**: Stable *CDKN1B* mRNA knockdown by two different shRNA constructs (sh1B-H3, sh1B-H4), shown using bar graph (mean ± SEM) in per^pos^ CTCs (BRx-82, BRx-142), compared with scrambled control. P-value calculated using two-tailed Student’s T-Test (**** p<0.0001). **4E:** *CDKN1B* knockdown suppresses regrowth of DTPs following pulse DTX treatment of per^pro^ CTCs (BRx-82, BRx-142). Infection of CTC cultures with either sh1B-H3 (red) or sh1B-H4 (green), followed by DTX exposure (10nM, 16 hrs) abolishes the regrowth observed with shScrambled control (blue). A representative graph, from at least two biological repeats, is shown. **4F:** *CDKN1B* knockdown efficiency following siRNA transfection of CTC cultures (BRx-142) is represented by bar graph (mean ± SEM). p-value was calculated using two-tailed Student’s T-Test (*** p<0.01). **4G:** Bar graphs (mean ± SEM) showing the quantification of mitotic (pHH3-positive) 4N and ≥8N BRx-142 CTCs, shown for untreated (NT, blue) and DTX-treated (red; 10nM, day 7) cultures, following transfection with either siCDKN1B or siControl. *CDKN1B* knockdown increases the number of mitotic >8N CTCs following pulse DTX, but not the number of mitotic 4N CTCs. P-value calculated using two-tailed Student’s T-Test (** p<0.01, ***p<0.001). n=3 biological triplicates.

To assess the functional contribution of *CDKN1B* to the appearance of DTX-induced persister cells, we knocked it down using siRNA in per^pro^ and per^def^ cultures (Supplementary Figure 4F), followed by 2 weeks of DTX exposure. *CDKN1B* knockdown significantly reduces the fraction of per^pro^ CTCs (BRx-68, BRx-82, BRx-142) surviving following DTX-treatment, compared with siRNA controls (1-E_max_ (relative viability) in si*CDKN1B* versus siControl were: BRx-68, 23.9 ± 0.15%versus 39.6 ± 1.35%, p<0.001; BRx-82, 2.3 ± 0.094%versus 9.1 ± 0.23%, p<0.00001; BRx-142, 23.2 ± 0.89%versus 37.1 ± 2.26%, p<0.01, Figure 4C). In contrast, *CDKN1B* knockdown in per^def^ CTCs (BRx-07, BRx-29, BRx-394) does not affect their survival following DTX treatment (Figure 4C). To test whether *CDKN1B* affects the ability of DTPs to resume proliferation following drug withdrawal, we used two different shRNA sequences to suppress its expression in per^pro^ CTCs (BRx-82, BRx-142; Figure 4D). Whereas scrambled shRNA-infected per^pro^ CTCs resume proliferation within 3-6 weeks after pulse DTX, *CDKN1B*-KD cells fail to regrow for up to 10 weeks (Figure 4E).

DTX-induced endomitosis, characterized by the appearance of mitotic polyploid (≥8N) cells, is more abundant in per^def^ CTCs, compared with per^pro^ cultures. Indeed, depletion of *CDKN1B* in per^pro^ CTCs (Figure 4F) leads to increased endomitosis following DTX exposure. This is evident in the higher fraction of ≥8N pHH3+ CTCs in *siCDKN1B* DTX-treated CTCs, compared to siControl CTCs (0.65 ± 0.03% vs 1.26 ± 0.1%, p<0.01; Figures 4G, right panel). No such differential increase mediated by *siCDKN1B* CTCs is observed among 4N cells (Figure 4G, left panel). Finally, the effect of si*CDKN1B* on the induction of drug-tolerant persisters appears to be inherent to treatment with mitotic inhibitors. Compared with control cells, *siCDKN1B* reduces the fraction of persisters following treatment with the microtubule polymerization inhibitors vincristine or the kinesin 5 inhibitor EMD534085, but it had no effect on cells treated with the DNA-damaging agent doxorubicin (Supplementary Figure 4G and 4H).

Taken together, depletion of CDKN1B in these patient-derived cultures leads to the acquisition of a per^def^ phenotype, with endomitosis leading to non-viable ≥8N cells.

### CDKN1B protein stabilization in DTPs is mediated through AKT phosphorylation

The increase in CDKN1B protein evident in per^pro^ CTCs following DTX exposure is not accompanied by an increase in *CDKN1B* mRNA (Supplementary Figure 5A), suggesting that it may involve post-transcriptional mechanisms. To determine whether protein stabilization is responsible for DTX-induced upregulation of CDKN1B protein, we used a chimeric reporter construct, mVenus-CDKN1B-K, in which a CMV promoter-driven nonfunctional CDKN1B is fused to the mVenus fluorescent^32^, providing a readily quantifiable measurement of post-transcriptional effects on CDKN1B protein (Methods). Expression of the reporter in per^pro^ CTCs (BRx-82, BRx-142) treated with pulse DTX shows a 5.3-fold increase in fluorescence, consistent with protein stabilization (Supplementary Figure 5B). Despite this level of protein stabilization, the ubiquitin E3 ligase SKP2, commonly implicated in the degradation of CDKN1B, is not altered in DTX-treated cells (supplementary Figure 5C). Since CDKN1B degradation is known to be modulated by its phosphorylation at multiple serine residues^33^, we tested whether DTX exposure leads to CDKN1B phosphorylation. We first generated a mutant mVenus-CDKN1B-K^-^, in which all 7 known CDKN1B phosphorylation targets are converted to alanine (mutant). DTX-mediated stabilization of the CDKN1B reporter is suppressed in this mutant construct, with reduced induction from 10.9-fold (WT) to 1.8-fold (mutant) in BRx-142 and from 3-fold (WT) to 1.2-fold (mutant) in BRx-82 CTCs (Figure 5A).

**Figure 5:**
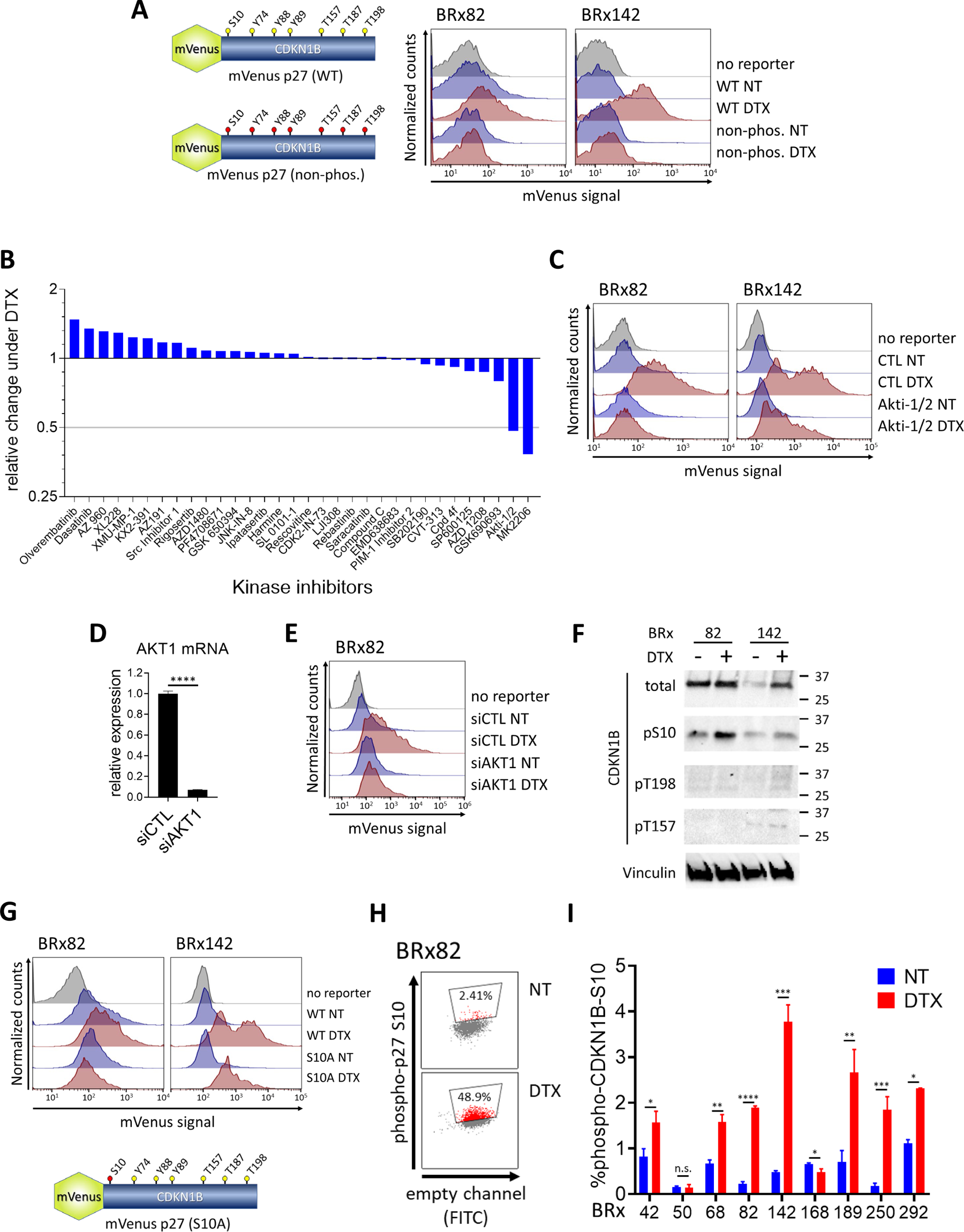
CDKN1B protein stabilization in persister cells is mediated through AKT phosphorylation. **5A:** Stabilization of mVenus-p27 reporter construct following DTX exposure. ***Left:*** schema of the reporter, fusing a fluorescent tag (mVenus) to the catalytically inactive CDKN1B coding sequence of *CDKN1B,* driven by the CMV promoter. Phosphorylation target residues are shown in the wild-type (WT) reporter, all of which are mutated to alanine in the mutant construct (non-phos.) ***Right:*** Increased mVenus fluorescence signal (flow cytometry) following treatment of CTC cultures (BRx-82, BRx-142) with DTX (10nM, 4 days), indicating stabilization of the reporter protein. Mutation of all 7 candidate phosphorylation target residues abrogates stabilization of the reporter. One of two biological repeats is shown. **5B**: Waterfall plot, showing a kinase inhibitor screen to identify candidates mediating stabilization of the mVenus-p27 reporter, following treatment of CTCs (BRx-142) with DTX (10nM, 4 days). Among 32 kinase inhibitors targeting 15 kinases previously implicated in stabilizing CDKN1B, two compounds targeting AKT (Akti-1/2 and MK2206) suppress DTX-mediated stabilization of CDKN1B. Bar graphs represent the ratio of the percent reporter-positive cells exposed to DTX along with inhibitor as a function of the percent reporter-positive cells exposed to DTX alone. **5C:** Suppression of CDKN1B reporter stabilization in CTC cultures (BRx-82, BRx-142) treated with DTX (10nM, 4 days) in the presence of the AKT inhibitor Akti-1/2 (100mM). mVenus fluorescence signal in live cells was measured using flow cytometry, with one of two biological repeats shown. **5D:** Knockdown of the *AKT1* isoform using siRNA transfection in CTC cultures (BRx-82) stably expressing the CDKN1B reporter. Bar graph (mean ± SEM), with p-value was calculated using two-tailed Student’s T-Test (**** p<0.0001 **5E:** Suppression of CDKN1B reporter stabilization in DTX-treated CTCs (BRx-82) following transfection with si*AKT1*, compared with siControl. Shown is one of two biological repeats. **5F:** Western blot analysis showing increased phosphorylation of one of the three AKT phosphorylation sites on native CDKN1B protein, serine-10 (S10), in CTC cultures (BRX-82, BRx-142) following treatment with DTX (10nM, 4 days). The other two AKT mediated phosphorylation sites (T198 and T157) are less evident by Western blotting . Total CDKN1B protein levels are shown (with and without DTX exposure), along with Vinculin expression (loading control). **5G:** Mutation of the *CDKN1B* S10 residue within the mVenus reporter is sufficient to reduce DTX-mediated protein stabilization. The position of the serine 10 residue (mutated to alanine) is shown in the schema (mVenus p27 (S10A). Mutation of S10 reduces CDKN1B stabilization in CTCs (BRx-82, BRx-142) following DTX exposure (10nM, 4 days), compared with wild type (WT) reporter. mVenus fluorescence signal in live cells measured using flow cytometry, with one of two biological repeats shown. **5H:** Flow cytometric analysis, showing increased subpopulation of CTC cultures (BRx-82) with detectable endogenous phospho-serine 10 (S10)-CDKN1B protein, following DTX exposure, compared with untreated cultures (NT). A representative scatter plot is shown. **5I:** Bar graph, showing quantification of the fraction of per^pro^ CTC cultures (BRx-42, BRx-50, BRx-68, BRx-82, BRx-142, BRx-168, BRx-189, BRx-292) with phosphorylated endogenous Serine 10 (S10)-CDKN1B. Percent S10-CDKN1B positive cells (mean ± SEM) following DTX exposure or untreated (NT) is shown, with p-value calculated using two-tailed Student’s T-Test (* p<0.05, ** p<0.01, *** p<0.001, **** p<0.0001).

To identify kinases that mediate the DTX-induced stabilization of CDKN1B, we tested the effect of 32 small molecule kinase inhibitors, targeting 15 different kinases implicated in CDKN1B phosphorylation in different cellular contexts^34–52^ (Supplementary table 3 and Figure 5B). Two AKT inhibitors, AKTi-1/2 and MK-2206, were most potent in suppressing CDKN1B stabilization following DTX exposure (Figure 5C). This effect is recapitulated by siRNA-mediated knockdown of *AKT1*, the predominant AKT isoform expressed in these cells (Supplementary Figure 5D), with a decrease in DTX-associated reporter induction from 4.6-fold to 1.6-fold (Figure 5D and 5E). AKT phosphorylates CDKN1B on three residues, serine-10 and threonine-157 and threonine-198^45, 47, 50^, leading us to test the phosphorylation of these residues by Western blot using phospho-specific antibodies. Phosphorylation of CDKN1B serine-10, but not threonine-157 or threonine-198, is evident by immunoblotting 4 days following DTX exposure (Figure 5F). Mutation of the serine-10 to an alanine residue within the mVenus-CDKN1B-K^-^ reporter is sufficient to abolish its accumulation following DTX exposure (Figure 5G). Finally, flow cytometry of multiple per^pro^ CTC lines using a phospho-serine-10 CDKN1B-specific antibody shows that DTX-treatment increases the subpopulation of cells with detectable phosphorylation at that residue in 7 of 9 cultures tested (BRx-42, BRx-50, BRx-68, BRx-82, BRx-142, BRx-168, BRx-189, BRx-250, BRx-292) from a median of 0.6% ± 0.13% (range 0.17% to 1.11%) in untreated cells to 2.24% ± 0.29% (range 1.57% to 3.77%) in DTX-treated CTCs (p-values range from 0.017 to 0.000042, Figure 5H and 5I). Together, these results indicate that the stabilization of CDKN1B following DTX exposure results from AKT1-mediated phosphorylation on serine-10.

## Discussion

In studying cultured CTCs derived from patients with advanced, heavily treated metastatic breast cancer, we describe a heritable phenotype, independent of clinical treatment history, whereby some CTC cultures are effectively killed by the mitotic inhibitor DTX, while others give rise to transient drug-resistant persisters and ultimately recover cell proliferation, resuming their parental drug susceptibility profile. We link the ability to generate DTX resistant persisters (per^pro^) to a more complete cell cycle arrest, which is lacking in cells that fail to regrow after a pulse of DTX (per^def^). Indeed, we find in per^def^ cells a higher population of highly polyploid (≥8N) mitotic cells following DTX exposure. In DTX treated per^pro^ CTC cultures, AKT1-mediated stabilization of the cell cycle inhibitor CDKN1B is required to prevent such endoreduplication, allowing for the emergence of the drug persister phenotype. Suppression of *CDKN1B* enhances cell killing by DTX and prevents the subsequent renewal of cell proliferation. The presence of this pathway in CTCs from some patients but not others is of particular interest, given the known cellular plasticity and loss of physiological cell cycle checkpoints that characterize advanced tumors and the common use of mitotic inhibitors to treat such refractory breast cancers (see Figure 6).

**Figure 6:**
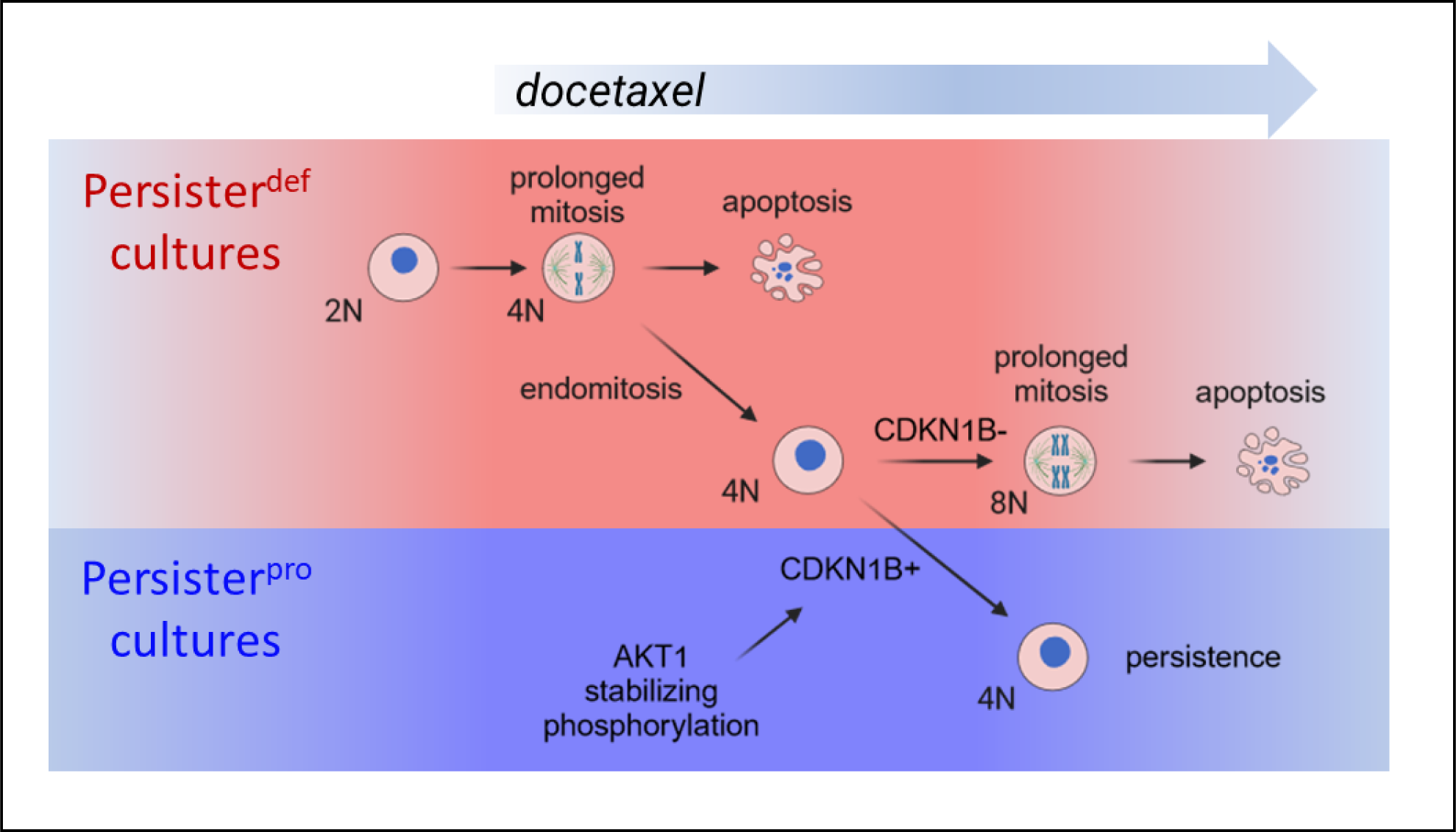
Proposed model showing CDKN1B-mediated restriction of endomitosis contributing to the viability of DTPs following DTX treatment. Advanced breast cancer cells from different patients, exemplified by *ex vivo* cultured CTCs, may contain subpopulations of cells susceptible to becoming drug-tolerant persisters (DTPs), *i.e*., per^pro^ cultures, or they may lack such cells, *i.e*., per^def^ cultures. Most cancer cells (both per^pro^ and per^neg^ cultures) exposed to the mitotic inhibitor DTX undergo prolonged mitosis followed by apoptosis, and a subset undergo endomitosis, with another prolonged mitosis at the 8N stage followed by cell death. Within per^pos^ CTC cultures, a subset of cells show AKT1-mediated stabilization of CDKN1B (p27), which restricts polyploidization. The resulting cell cycle arrested but viable (4N) cells constitute DTPs. In the absence of DTX, these DTPs eventually reenter the cell cycle (2N), with drug sensitivity comparable to that of their parental cells.

In normal cells, CDKN1B plays a critical role in suppressing G1/S transition by inhibiting cyclin D-CDK4/6 and cyclin E-CDK2 complexes^53, 54^. Compared with its close family member CDKN1A, which is commonly implicated in DNA damage-induced checkpoints, CDKN1B appears to mediate developmentally regulated cell cycle regulation and maintain long-term quiescence following normal proliferation^54^. In the setting of advanced, treatment-refractory breast cancer, CDKN1B, rather than CDKN1A, mediates a drug-induced mitotic checkpoint that appears critical to allow cells to survive acute drug-induced mitotic injury, setting the stage for the emergence of DTPs. While targeting CDKN1B might enhance DTX efficacy in advanced breast cancer, it encodes an intrinsically unstructured protein which is thought to be a poor drug target^55^. In this context, our finding that AKT mediates CDKN1B stabilization following DTX exposure might also offer a therapeutic opportunity. AKT modulates multiple signaling pathways implicated in both cell survival and proliferation, making it a complex drug target with potentially opposing functional consequences. Moreover, mTOR inhibition itself may induce a persister state in pancreatic cancer cells^54^. Whereas direct knockdown of *CDKN1B* suppresses the DTX-induced DTP phenomenon in our CTC model system, we were unable to identify a drug sequencing regimen in which the combination of DTX and AKT inhibitor prevents regrowth of DTPs without massively increased cell toxicity. Of note, while multiple AKT inhibitors have been tested in clinical trials, their combination with taxanes in the treatment of advanced breast cancer have yielded limited results^56, 57^, possibly due to lack of patient stratification. Thus, any role for AKT inhibitors in combinational therapies to prevent drug resistance in the clinical setting remains to be defined.

Our results also shed light on the mechanism of DTX-induced cell death in advanced, refractory breast cancer. Taxanes are thought to induce cell death through two potential mechanisms^58^. In some cells, cell death results from prolonged mitotic arrest, linked to the degradation of BH3 family anti-apoptotic proteins, including MCL1, BCL2, and BCLXL^16–18, 58^, thereby tilting the mitochondrial balance toward apoptosis. Alternatively, taxane-treated cells may undergo mitotic slippage^59^, whereby the nuclear membrane reassembles without cytokinesis, producing one or more cycles of endoreduplication, endomitosis and aneuploidy^59–61^. Indeed, the primary distinction between per^pro^ and per^def^ cultured CTCs appears to be the prevalence of cells with 8N DNA content expressing the mitotic marker pHH3, and it is this second endoreduplication mechanism that is dependent on CDKN1B. By suppressing this pathway, CDKN1B appears to enable the emergence of DTPs.

Transient drug-tolerant persister populations are increasingly appreciated as an intermediate state that may precede the acquisition of stable cancer drug resistance through fixed mutational mechanisms^1^. Once established, persister states appear to share common features of cellular quiescence, independent of the initiating therapeutic intervention^5^. However, the early steps in the initiation of a drug-tolerant cell state may involve drug-specific pathways that avoid immediate cell death, as demonstrated in our study of DTX-induced DTPs within breast CTC cultures. Understanding and ultimately targeting such early drug pathway-specific mechanisms may enable the prevention of the DTP phenomenon to enhance cancer drug efficacy. The transient DTX resistance phenomenon in CTCs, together with the DTX sensitivity of CTCs drawn from some patients whose tumors had previously been thought to have acquired resistance to taxanes, may also help explain cases in which patients derive benefit from repeated intermittent treatment with mitotic inhibitors.

## Methods

### CTC isolation and culture

Patients diagnosed with metastatic breast cancer consented to de-identified blood collection in accordance with the protocol approved by the institutional review board (DF/HCC 05-300). Enrolled patients had undergone multiple courses of therapy, commonly applied in advanced HR+ breast cancer. Patient-matched primary and metastatic tumor specimens were collected following the institutional review board approved protocol (2002-P-002059).

Isolation of CTCs from fresh whole blood was carried out by depleting leukocytes using the microfluidic CTC-iChip, as previously detailed^27^. In brief, whole blood samples were incubated with biotinylated antibodies against CD45 (R&D Systems, clone 2D1), CD66b (AbD Serotec, clone 80H3), and CD16 (BD, clone 3G8). Subsequent incubation with Dynabeads MyOne Streptavidin T1 (Invitrogen) facilitated magnetic labeling of white blood cells. This mixture underwent processing through the CTC-iChip to deplete leukocytes, enabling collection of untagged, viable CTCs.

CTC cultures were grown in suspension in ultra-low attachment plates (Corning) in tumor sphere medium - RPMI-1640, EGF (20 ng/ml), bFGF (20 ng/ml), 1X B27, 1X antibiotic/antimycotic (Life Technologies) under hypoxic (4% O_2_) conditions. CTC lines were routinely checked for mycoplasma, using a mycoplasma detection kit (MycoAlert, Lonza). For bulk lysate-based analyses (RNA sequencing and western blot), live cells were separated from dead cells using Ficoll-Paque Plus (Cytiva) enrichment by centrifuging at 400×g for 5 minutes at room temperature and removing the top layer and washing it with PBS.

### Fluorescence-activated cell sorting (FACS)

Prior to staining, cells were labeled with LIVE/DEAD fixable far red dead cell stain kit (Invitrogen), diluted in PBS for 5 minutes at room temperature. Cells were fixed with Cytofix/Cytoperm Fixation/Permeabilization Kit (BD) for 10 minutes at room temperature followed by 10 minutes incubation with Perm/Wash buffer. Intracellular stain with antibodies was performed in Perm/Wash buffer. Antibodies for immunostaining were: Phospho-Histone H3 Serine 10 PE conjugated (Cell Signaling), mouse anti CDKN1B (Novus), mouse IgG1 control (Novus), rabbit anti phospho-CDKN1B Serine 10 (Abcam), rabbit IgG control (Cell Signaling). Cells were incubated with the different antibodies for 30 minutes on ice. Secondary antibodies were goat anti mouse Alexa-546 and goat anti rabbit Alexa-555 (Invitrogen). DNA was stained with Hoechst 33342 (Thermo Scientific) at 20µM, for 5 minutes in Perm/Wash buffer at room temperature. Flow based measurements of endogenous mVenus fluorescence was measured on non-fixed cells labeled with LIVE/DEAD stain. For single cell RNA sequencing, cells were sorted with a LIVE/DEAD far red stain using a MA900 Multi-Application Cell Sorter (SONY). Live single cells were sorted into 96 wells plates containing 10µl of TCL lysis buffer (Qiagen 1031576), supplemented with 1% β-mercaptoethanol.

### Drug treatments, cell viability assays and drug screens

Drug kill curves were performed using Cell Titer Glo 2.0 (Promega) on a SpectraMax M5 plate reader (Molecular Devices). Chemical compounds used in this work were Docetaxel (Selleck), Doxorubicin (Sigma Aldrich), EMD534085 (Medchem Express), Vincristine (Selleck), AKTi-1/2 (Tocris), MK-2206 (VWR). A custom library of kinase inhibitors was purchased from TargetMol. Cell counts for long term viability assays were carried using CellDrop (Denovix) on brightfield mode. Jpeg image files were exported, and manual counting was performed using the ImageJ software (https://imagej-nih-gov.ezp-prod1.hul.harvard.edu/ij/).

### Single cell and bulk RNA sequencing

For single cell RNA sequencing, libraries from single cell lysates were generated as previously described^62^ using the Smart-Seq2 protocol^63^. 96-well plates containing cell lysates were thawed on ice, spun down at 1500 rpm for 30s, and mixed with Agencourt RNAClean XP SPRI beads (Beckman Coulter) for RNA purification. Purified RNA was resuspended in 4μl of Mix-1, denatured at 72°C for 3 min and transferred to ice for 1 min before 7μl of Mix-2 was added. Reverse transcription was carried out at 50°C for 90 min, followed by 5 min incubation at 85°C. 14μl of Mix-3 was added in each well and the whole-transcriptome amplification step was performed at 98°C for 3 min, followed by 21 cycles at (98°C for 15 s, 67°C for 20 s and 72°C for 6 min), and final extension at 72°C for 5min. cDNA was then purified with Agencourt AMPureXP SPRI beads (Beckman Coulter), to remove all primer dimer residues. cDNA concentration measurements were performed using the Qubit dsDNA high sensitivity assay kit on the Synergy H1 Hybrid Microplate Reader (BioTek). Libraries were generated using the Nextera XT Library Prep kit (Illumina) with custom indexing adapters^64^ in a 384-well PCR plate, followed by a cleanup step to remove residual primer dimers. Combined libraries from different cells were then sequenced on a NextSeq 500 sequencer (Illumina), using paired-end 38-base reads.

### Bulk RNA-seq data analysis

Trimmomatic was used to crop reads lengths to 50 nucleotides, and to remove the TruSeq3-PE-2 Illumina adapters. The paired-end reads were then aligned using tophat2 and bowtie1 with the no-novel-juncs argument set and with transcriptome defined by the hg19 genes.gtf table from http://genome.ucsc.edu. Reads that did not align or aligned to multiple locations were discarded. The number of reads aligning to each gene was then determined using htseq-count. The read count for each gene was divided by the total counts assigned to all genes and multiplied by one million to form the reads per million (RPM).

### Single-cell RNA-seq data and UMAP analyses

Raw fastq reads generated from the sequencer were first cleaned using TrimGalore (v0.4.3) (https://github.com/FelixKrueger/TrimGalore) to remove the adapter-polluted reads and reads with low sequencing quality. Cleaned reads were aligned to the human genome (hg19) using Tophat (v2.1.1)^65^. PCR duplicates were further removed using samtools (v1.3.1)^66^, gene counts were computed using HTseq (v0.6.1)^67^. For UMSP anslysis, we discarded samples with fewer than 400,000 alignments or with less than 30% of genes having at least one aligning read. We then used the prcomp function of the stats package of R to compute the first 15 principal components of the RPM (Reads Per Million) matrix. We then ran the umap function of version 0.2.10.0 of the umap package of R <arXiv:2802.03426> with the first 15 principal components as input. Every sample in this analysis has a treatment status of either no treatment (NT) or docetaxel (DTX). For each sample we determine whether its nearest neighbor in the 2-dimensional output of umap has the same treatment status as the sample. This gives us a measure of how segregated by treatment status the samples are. To determine whether the degree of segregation is more than would be expected by chance, we performed a permutation test in which we randomly permuted the treatment statuses of the samples and computed the resulting degree of segregation. Doing this for a large number (100,000) of random permutations yields an empirical null distribution. The portion of the empirical null distribution that is greater than the actual degree of segregation is the p-value determined by the permutation test. Computing metascores for E2F and proliferation was performed using the HALLMARK_E2F_TARGETS gene set and the WHITFIELD_CELL_CYCLE_LITERATURE gene set in version 6.0 of the MSigDB gene set database [https://www.gsea-msigdb.org/gsea/msigdb], we defined a gene expression metascore as the mean of the log10(RPM + 1) values for the genes in the gene set.

### Phosphoproteomics

Frozen cell pellets were lysed, and proteins were reduced using DTT and alkylated with iodoacetamide, then precipitated using the MeOH/CHCl3 protocol and digested using LysC and trypsin, followed by phosphopeptide enrichment as previously described^30^. For each sample 2.5 mg of peptides were subjected to phosphopeptide enrichment on TiO2 beads (GL Sciences, Japan). Phosphopeptides were labeled with TMT10plex reagents (Thermo Fisher Scientific), pooled, and were fractionated into 24 fractions using basic pH reverse phase chromatography essentially as described previously^68^. Samples were dried, re-suspended in 5% ACN/5% formic acid, and analyzed in 3-hour runs via LC-M2/MS3 on an Orbitrap FusionLumos mass spectrometer using the Simultaneous Precursor Selection (SPS) supported MS3 method^69, 70^ essentially as described^71^. Two MS2 spectra were acquired for each peptide using CID and HCD fragmentation as described^72^ and the gained MS2 spectra were assigned using a SEQUEST-based in-house built proteomics analysis platform^73^ allowing phosphorylation of serine, threonine, and tyrosine residues as a variable modification. The Ascore algorithm was used to evaluate the correct assignment of phosphorylation within the peptide sequence^74^ . Based on the target-decoy database search strategy^75^ and employing linear discriminant analysis and posterior error histogram sorting, peptide and protein assignments were filtered to false discovery rate (FDR) of ˂ 1%^73^. Peptides with sequences that were contained in more than one protein sequence from the UniProt database (2014) were assigned to the protein with most matching peptides^73^. Only MS3 with an average signal-to-noise value of larger than 40 per reporter ion as well as with an isolation specificity^70^ of larger than 0.75 were considered for quantification. A two-step normalization of the protein TMT-intensities was performed by first normalizing the phospho-peptide intensities over all acquired TMT channels for each protein based on the median average peptide intensity calculated for all peptides. To correct for slight mixing errors of the peptide mixture from each sample a median of the normalized intensities was calculated from all peptide intensities in each TMT channel and the peptide intensities were normalized to the median value of these median intensities.

### Proteomic analyses

50 µg of the peptides resulting after tryptic digest as described above were subsequently labeled using TMT-10plex reagents (Thermo Scientific) according to manufacturer’s instructions. Labeled samples were combined and fractionated using a basic reversed phase HPLC^68^. The resulting fractions were analyzed in a 3h reversed phase LC-MS2/MS3 run on an Orbitrap Fusion. MS3 isolation for quantification used Simultaneous Precursor Selection (SPS) as previously described^69–71^. Proteins were identified based on MS2 spectra using the sequest algorithm searching against a human data base (uniprot 2014)^76^ using an in house-built platform^73^. Search strategy included a target-decoy database-based search to filter against a false-discovery rate (FDR) of protein identifications of less than 1%^75^. For quantification only MS3 with an average signal-to-noise value of larger than 40 per reporter ion as well as with an isolation specificity^70^ of larger than 0.75 were considered and a two-step normalization as described above was performed. Protein intensities are normalized using an analogous methodology as the phospho-proteomics.

### Lenti-vectors cloning and production

For the cloning of mVenus-CDKN1B-K^-^ reporter, a DNA sequence of 1332 base pairs encompassing the gene for mVenus in frame with the CDKN1B gene and flanked by *BamHI* and *XhoI* inside the subcloning vector pUC-GW-Amp was ordered from GENEWIZ. The CDKN1B in this reporter is inactivated by mutations to alanine in two of its phenyl-alanine residues (F62A-F64A), rendering it unable to bind the CDK proteins^32^. This fragment was subcloned by restriction (*BamHI* and *XhoI*) and ligation into the lenti-vector expression backbone plasmid pLenti-CMV-Blast-empty-(w263-1) (Addgene). Sequence integrity was validated with Sanger sequencing. To produce the CDKN1B non-phosphorylation mutant reporter, a similar sequence was ordered from GENWIZ with 7 of its phosphoresidues mutated in alanine. The CDKN1B Serine 10 to alanine mutant reporter was cloned using the Quikchange II site-directed mutagenesis kit (Agilent).

Lenti-vectors were prepared in 293T cells using X-tremeGene HP DNA transfection reagent (Roche), with psPAX2 and pMD2.G (Addgene) as enzyme and packaging carriers. Vector particle containing supernatants were collected at 72 hours past transfection. Transduction was carried through an overnight incubation of lenti-vector supernatants with 8 µg/ml Polybrene (Santa Cruz). Transduced cells were selected for with 10 µg/ml Blasticidin S HCL (Gibco).

### siRNA and shRNA knockdowns

Targeted and transient disruption of *CDKN1B* and *AKT1* was performed using siRNA duplexes (ON_TARGET plus smart pool siRNA Horizon Discovery). A non-targeting siRNA pool was used as control. siRNA duplexes were introduced into the BRx cultures using Lipofectamine RNAiMAX transfection reagent (Thermo Fisher). Growth media were replaced at 16 hours past transfection. CDKN1B and scrambled control shRNA in the pLKO.1 lentiviral backbone vector were purchased from the TRC shRNA library (Broad Institute). Selection for shRNA transduced cells was performed using 2 µg/ml Puromycin DiHCL (Thermo Scientific).

### RNA isolation, reverse transcription and quantitative PCR

RNA was isolated from live cells using the Quick-RNA Microprep Kit (Zymo). Quantification of RNA concentration was performed using NanoDrop One (Thermo Scientific). cDNA from 300ng of each sample’s RNA was synthesized using qScript cDNA Synthesis Kit, (QuantaBio). qPCR was performed using Taqman probe assays (Thermo Scientific) and using Taqman 2X Universal PCR Master Mix (Applied Biosystems). qPCR was performed on CFX Opus 384 thermal cycler (Bio-Rad). Analysis of qPCR was carried using the CFX Maestro software (Bio-Rad).

### Western blot analyses

Cells were lysed using Laemmli buffer (Sigma) supplied with Halt Protease and Phosphatase Inhibitor Cocktail (Thermo Scientific). Lysed cells were boiled at 95°C for 5 minutes. Protein concentrations were quantified using DC Protein Assay (Bio-Rad) and compared to an Albumin Standard (Thermo Scientific). 20µg of protein was loaded onto Mini-Protean TGX 4-20% gradient gels (Bio-Rad). Gels were run on a Mini-PROTEAN Tetra Vertical Electrophoresis Cell (Bio-Rad) with Tris-Glycine-SDS Running Buffer (Boston BioProducts) and with Dual Color Precision Plus Protein Standards (Bio-Rad). Proteins were then transferred to a 0.2µm Nitrocellulose Membrane (Bio-Rad) through wet transfer on the Mini-PROTEAN Tetra Vertical Electrophoresis Cell (Bio-Rad) with 10% Methanol added to the Transfer Buffer (Boston BioProducts). Membranes were blocked for 1 hour with 5% Non-Fat Powdered Milk (Boston BioProducts). Primary antibodies were incubated overnight. Washes were performed with PBS-T (Boston BioProducts). HRP conjugated secondary goat anti-mouse and goat anti-rabbit antibodiesxs (Bio-Rad) at 1:10,000 dilution. Immobilon Crescendo Western HRP substrate (1 minute in room temperature) was applied to membrane for chemiluminescence imaging. ChemiDocMP (Bio-Rad) was used to image the chemiluminescent signal. Primary antibodies used for western blot were rabbit mAb anti p27 Kip1 (Cat# 3686, Cell signaling), mouse anti-Vinculin (Cat# MAB3574, Millipore), rabbit anti-p27 Kip1 phospho-Serine 10 (Cat# ab62364, Abcam), rabbit anti-p27 KIP 1 phospho Threonine 198 antibody (Cat# ab64949, Abcam), rabbit anti Human Phospho-p27/Kip1 Threonine 157 (Cat# AF1555, R&D Systems).

### Statistical analysis

Unless noted otherwise, data were analyzed using a paired two-tail Student t test (for two-group comparisons) and two-way ANOVA for repeated measures. Data were processed using Microsoft Excel 365. Graphs were generated using GraphPad Prism 9 and Excel. Error bars represent 1 SEM.

## References

1. Sharma, S.V. et al. A chromatin-mediated reversible drug-tolerant state in cancer cell subpopulations. Cell 141, 69–80 (2010).

2. Dhimolea, E. et al. An Embryonic Diapause-like Adaptation with Suppressed Myc Activity Enables Tumor Treatment Persistence. Cancer Cell 39, 240–256 e211 (2021).

3. Echeverria, G.V. et al. Resistance to neoadjuvant chemotherapy in triple-negative breast cancer mediated by a reversible drug-tolerant state. Sci Transl Med 11 (2019).

4. Hangauer, M.J. et al. Drug-tolerant persister cancer cells are vulnerable to GPX4 inhibition. Nature 551, 247–250 (2017).

5. Recasens, A. & Munoz, L. Targeting Cancer Cell Dormancy. Trends Pharmacol Sci 40, 128–141 (2019).

6. Oren, Y. et al. Cycling cancer persister cells arise from lineages with distinct programs. Nature 596, 576–582 (2021).

7. Cabanos, H.F. & Hata, A.N. Emerging Insights into Targeted Therapy-Tolerant Persister Cells in Cancer. Cancers (Basel) 13 (2021).

8. Guler, G.D. et al. Repression of Stress-Induced LINE-1 Expression Protects Cancer Cell Subpopulations from Lethal Drug Exposure. Cancer Cell 32, 221–237 e213 (2017).

9. Viswanathan, V.S. et al. Dependency of a therapy-resistant state of cancer cells on a lipid peroxidase pathway. Nature 547, 453–457 (2017).

10. Kurppa, K.J. et al. Treatment-Induced Tumor Dormancy through YAP-Mediated Transcriptional Reprogramming of the Apoptotic Pathway. Cancer Cell 37, 104–122 e112 (2020).

11. Jenks, A.D. et al. Primary Cilia Mediate Diverse Kinase Inhibitor Resistance Mechanisms in Cancer. Cell Rep 23, 3042–3055 (2018).

12. Rehman, S.K. et al. Colorectal Cancer Cells Enter a Diapause-like DTP State to Survive Chemotherapy. Cell 184, 226–242 e221 (2021).

13. Wang, T.H., Wang, H.S. & Soong, Y.K. Paclitaxel-induced cell death: where the cell cycle and apoptosis come together. Cancer 88, 2619–2628 (2000).

14. Bumbaca, B. & Li, W. Taxane resistance in castration-resistant prostate cancer: mechanisms and therapeutic strategies. Acta Pharm Sin B 8, 518–529 (2018).

15. Tan, M.H. et al. Specific kinesin expression profiles associated with taxane resistance in basal-like breast cancer. Breast Cancer Res Treat 131, 849–858 (2012).

16. Wertz, I.E. et al. Sensitivity to antitubulin chemotherapeutics is regulated by MCL1 and FBW7. Nature 471, 110–114 (2011).

17. Gazitt, Y. et al. Bcl-2 overexpression is associated with resistance to paclitaxel, but not gemcitabine, in multiple myeloma cells. Int J Oncol 13, 839–848 (1998).

18. Liu, J.R. et al. Bcl-xL is expressed in ovarian carcinoma and modulates chemotherapy-induced apoptosis. Gynecol Oncol 70, 398–403 (1998).

19. Strobel, T., Swanson, L., Korsmeyer, S. & Cannistra, S.A. BAX enhances paclitaxel-induced apoptosis through a p53-independent pathway. Proc Natl Acad Sci U S A 93, 14094–14099 (1996).

20. Gradishar, W.J. Taxanes for the treatment of metastatic breast cancer. Breast Cancer (Auckl) 6, 159–171 (2012).

21. Miyamoto, D.T., Ting, D.T., Toner, M., Maheswaran, S. & Haber, D.A. Single-Cell Analysis of Circulating Tumor Cells as a Window into Tumor Heterogeneity. Cold Spring Harb Symp Quant Biol 81, 269–274 (2016).

22. Yu, M. et al. Cancer therapy. Ex vivo culture of circulating breast tumor cells for individualized testing of drug susceptibility. Science 345, 216–220 (2014).

23. Yu, M. et al. Circulating breast tumor cells exhibit dynamic changes in epithelial and mesenchymal composition. Science 339, 580–584 (2013).

24. Chemi, F. et al. Early Dissemination of Circulating Tumor Cells: Biological and Clinical Insights. Front Oncol 11, 672195 (2021).

25. Pantel, K. & Alix-Panabieres, C. Crucial roles of circulating tumor cells in the metastatic cascade and tumor immune escape: biology and clinical translation. J Immunother Cancer 10 (2022).

26. Cortes-Hernandez, L.E., Eslami, S.Z., Pantel, K. & Alix-Panabieres, C. Circulating Tumor Cells: From Basic to Translational Research. Clin Chem 70, 81–89 (2024).

27. Ozkumur, E. et al. Inertial focusing for tumor antigen-dependent and -independent sorting of rare circulating tumor cells. Sci Transl Med 5, 179ra147 (2013).

28. Ebright, R.Y. et al. Deregulation of ribosomal protein expression and translation promotes breast cancer metastasis. Science 367, 1468–1473 (2020).

29. Jordan, N.V. et al. HER2 expression identifies dynamic functional states within circulating breast cancer cells. Nature 537, 102–106 (2016).

30. Kreuzer, J., Edwards, A. & Haas, W. Multiplexed quantitative phosphoproteomics of cell line and tissue samples. Methods Enzymol 626, 41–65 (2019).

31. Johnson, J. et al. Targeting the RB-E2F pathway in breast cancer. Oncogene 35, 4829–4835 (2016).

32. Oki, T. et al. A novel cell-cycle-indicator, mVenus-p27K-, identifies quiescent cells and visualizes G0-G1 transition. Sci Rep 4, 4012 (2014).

33. Razavipour, S.F., Harikumar, K.B. & Slingerland, J.M. p27 as a Transcriptional Regulator: New Roles in Development and Cancer. Cancer Res 80, 3451–3458 (2020).

34. Boehm, M. et al. A growth factor-dependent nuclear kinase phosphorylates p27(Kip1) and regulates cell cycle progression. EMBO J 21, 3390–3401 (2002).

35. Chu, I. et al. p27 phosphorylation by Src regulates inhibition of cyclin E-Cdk2. Cell 128, 281–294 (2007).

36. Deng, X., Mercer, S.E., Shah, S., Ewton, D.Z. & Friedman, E. The cyclin-dependent kinase inhibitor p27Kip1 is stabilized in G(0) by Mirk/dyrk1B kinase. J Biol Chem 279, 22498–22504 (2004).

37. Fujita, N., Sato, S. & Tsuruo, T. Phosphorylation of p27Kip1 at threonine 198 by p90 ribosomal protein S6 kinases promotes its binding to 14-3-3 and cytoplasmic localization. J Biol Chem 278, 49254–49260 (2003).

38. Grimmler, M. et al. Cdk-inhibitory activity and stability of p27Kip1 are directly regulated by oncogenic tyrosine kinases. Cell 128, 269–280 (2007).

39. Hong, F. et al. mTOR-raptor binds and activates SGK1 to regulate p27 phosphorylation. Mol Cell 30, 701–711 (2008).

40. Jakel, H., Weinl, C. & Hengst, L. Phosphorylation of p27Kip1 by JAK2 directly links cytokine receptor signaling to cell cycle control. Oncogene 30, 3502–3512 (2011).

41. Kim, H. et al. JNK signaling activity regulates cell-cell adhesions via TM4SF5-mediated p27(Kip1) phosphorylation. Cancer Lett 314, 198–205 (2012).

42. Kossatz, U. et al. C-terminal phosphorylation controls the stability and function of p27kip1. EMBO J 25, 5159–5170 (2006).

43. Lee, J.G. & Kay, E.P. Two populations of p27 use differential kinetics to phosphorylate Ser-10 and Thr-187 via phosphatidylinositol 3-Kinase in response to fibroblast growth factor-2 stimulation. J Biol Chem 282, 6444–6454 (2007).

44. Liang, J. et al. The energy sensing LKB1-AMPK pathway regulates p27(kip1) phosphorylation mediating the decision to enter autophagy or apoptosis. Nat Cell Biol 9, 218–224 (2007).

45. Liang, J. et al. PKB/Akt phosphorylates p27, impairs nuclear import of p27 and opposes p27-mediated G1 arrest. Nat Med 8, 1153–1160 (2002).

46. Morishita, D., Katayama, R., Sekimizu, K., Tsuruo, T. & Fujita, N. Pim kinases promote cell cycle progression by phosphorylating and down-regulating p27Kip1 at the transcriptional and posttranscriptional levels. Cancer Res 68, 5076–5085 (2008).

47. Nacusi, L.P. & Sheaff, R.J. Akt1 sequentially phosphorylates p27kip1 within a conserved but non-canonical region. Cell Div 1, 11 (2006).

48. Patel, P. et al. Brk/Protein tyrosine kinase 6 phosphorylates p27KIP1, regulating the activity of cyclin D-cyclin-dependent kinase 4. Mol Cell Biol 35, 1506–1522 (2015).

49. Rousseau, S. et al. CXCL12 and C5a trigger cell migration via a PAK1/2-p38alpha MAPK-MAPKAP-K2-HSP27 pathway. Cell Signal 18, 1897–1905 (2006).

50. Shin, I. et al. PKB/Akt mediates cell-cycle progression by phosphorylation of p27(Kip1) at threonine 157 and modulation of its cellular localization. Nat Med 8, 1145–1152 (2002).

51. Viglietto, G. et al. Cytoplasmic relocalization and inhibition of the cyclin-dependent kinase inhibitor p27(Kip1) by PKB/Akt-mediated phosphorylation in breast cancer. Nat Med 8, 1136–1144 (2002).

52. Zheng, Y.L. et al. Phosphorylation of p27Kip1 at Thr187 by cyclin-dependent kinase 5 modulates neural stem cell differentiation. Mol Biol Cell 21, 3601–3614 (2010).

53. Chu, I.M., Hengst, L. & Slingerland, J.M. The Cdk inhibitor p27 in human cancer: prognostic potential and relevance to anticancer therapy. Nat Rev Cancer 8, 253–267 (2008).

54. Abbastabar, M. et al. Multiple functions of p27 in cell cycle, apoptosis, epigenetic modification and transcriptional regulation for the control of cell growth: A double-edged sword protein. DNA Repair (Amst) 69, 63–72 (2018).

55. Bencivenga, D., Stampone, E., Roberti, D., Della Ragione, F. & Borriello, A. p27(Kip1), an Intrinsically Unstructured Protein with Scaffold Properties. Cells 10 (2021).

56. Cerma, K. et al. Targeting PI3K/AKT/mTOR Pathway in Breast Cancer: From Biology to Clinical Challenges. Biomedicines 11 (2023).

57. Martorana, F. et al. AKT Inhibitors: New Weapons in the Fight Against Breast Cancer? Front Pharmacol 12, 662232 (2021).

58. Mosca, L., Ilari, A., Fazi, F., Assaraf, Y.G. & Colotti, G. Taxanes in cancer treatment: Activity, chemoresistance and its overcoming. Drug Resist Updat 54, 100742 (2021).

59. Gascoigne, K.E. & Taylor, S.S. Cancer cells display profound intra- and interline variation following prolonged exposure to antimitotic drugs. Cancer Cell 14, 111–122 (2008).

60. Bolgioni, A.F., Vittoria, M.A. & Ganem, N.J. Long-term Live-cell Imaging to Assess Cell Fate in Response to Paclitaxel. J Vis Exp (2018).

61. Ganem, N.J. & Pellman, D. Linking abnormal mitosis to the acquisition of DNA damage. J Cell Biol 199, 871–881 (2012).

62. Sade-Feldman, M. et al. Defining T Cell States Associated with Response to Checkpoint Immunotherapy in Melanoma. Cell 176, 404 (2019).

63. Picelli, S. et al. Smart-seq2 for sensitive full-length transcriptome profiling in single cells. Nat Methods 10, 1096–1098 (2013).

64. Villani, A.C. & Shekhar, K. Single-Cell RNA Sequencing of Human T Cells. Methods Mol Biol 1514, 203–239 (2017).

65. Trapnell, C. et al. Differential gene and transcript expression analysis of RNA-seq experiments with TopHat and Cufflinks. Nat Protoc 7, 562–578 (2012).

66. Li, H. et al. The Sequence Alignment/Map format and SAMtools. Bioinformatics 25, 2078–2079 (2009).

67. Anders, S., Pyl, P.T. & Huber, W. HTSeq--a Python framework to work with high-throughput sequencing data. Bioinformatics 31, 166–169 (2015).

68. Edwards, A. & Haas, W. Multiplexed Quantitative Proteomics for High-Throughput Comprehensive Proteome Comparisons of Human Cell Lines. Methods Mol Biol 1394, 1–13 (2016).

69. McAlister, G.C. et al. MultiNotch MS3 enables accurate, sensitive, and multiplexed detection of differential expression across cancer cell line proteomes. Anal Chem 86, 7150–7158 (2014).

70. Ting, L., Rad, R., Gygi, S.P. & Haas, W. MS3 eliminates ratio distortion in isobaric multiplexed quantitative proteomics. Nat Methods 8, 937–940 (2011).

71. Erickson, B.K. et al. Evaluating multiplexed quantitative phosphopeptide analysis on a hybrid quadrupole mass filter/linear ion trap/orbitrap mass spectrometer. Anal Chem 87, 1241–1249 (2015).

72. Lyons, J. et al. Integrated in vivo multiomics analysis identifies p21-activated kinase signaling as a driver of colitis. Sci Signal 11 (2018).

73. Huttlin, E.L. et al. A tissue-specific atlas of mouse protein phosphorylation and expression. Cell 143, 1174–1189 (2010).

74. Beausoleil, S.A., Villen, J., Gerber, S.A., Rush, J. & Gygi, S.P. A probability-based approach for high-throughput protein phosphorylation analysis and site localization. Nat Biotechnol 24, 1285–1292 (2006).

75. Elias, J.E. & Gygi, S.P. Target-decoy search strategy for increased confidence in large-scale protein identifications by mass spectrometry. Nat Methods 4, 207–214 (2007).

76. Eng, J.K., McCormack, A.L. & Yates, J.R. An approach to correlate tandem mass spectral data of peptides with amino acid sequences in a protein database. J Am Soc Mass Spectrom 5, 976–989 (1994).

77. Ebright, R.Y. et al. HIF1A signaling selectively supports proliferation of breast cancer in the brain. Nat Commun 11, 6311 (2020).

78. Brett, J.O., et al. A Gene Panel Associated With Abemaciclib Utility in ESR1-Mutated Breast Cancer After Prior Cyclin-Dependent Kinase 4/6-Inhibitor Progression. JCO Precis Oncol 7, e2200532 (2023).

79. Whitfield, M.L. et al. Identification of genes periodically expressed in the human cell cycle and their expression in tumors. Mol Biol Cell 13, 1977–2000 (2002).

