## Supplementary Figures and Tables for "*CDKN1B* (p27^kip1^) enhances drug tolerant persister CTCs by restricting polyploidy following mitotic inhibitors": Horwitz et al - Supplementary Figures.pdf

### Supplementary Figure 1

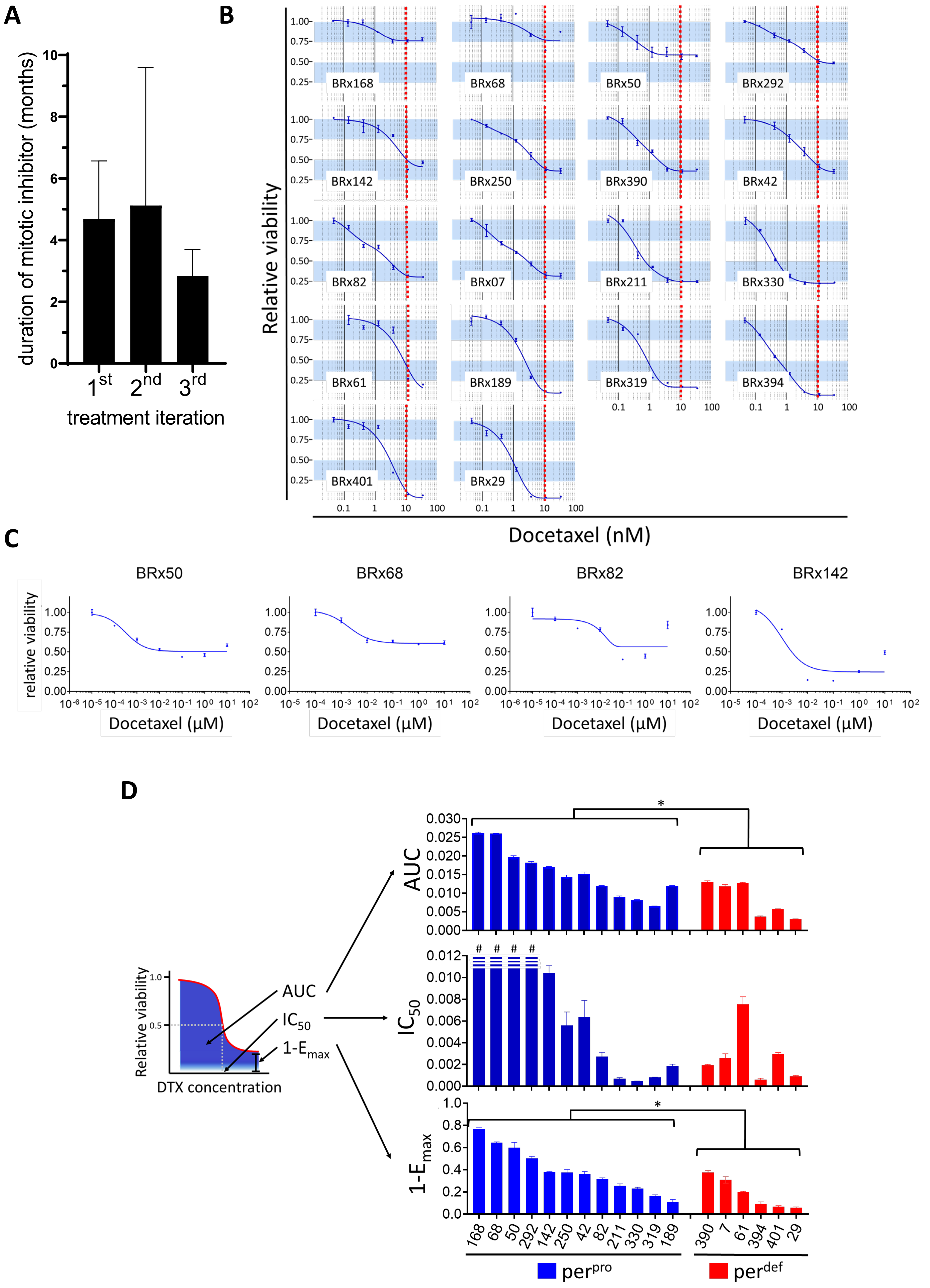

### Supplementary Figure 2

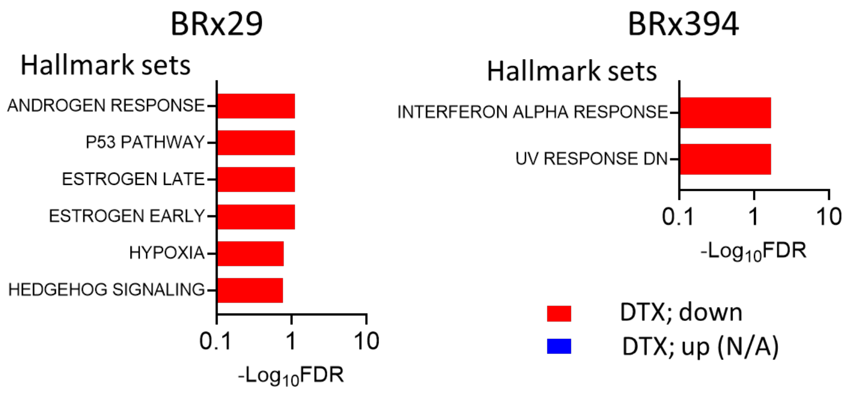

### Supplementary Figure 3

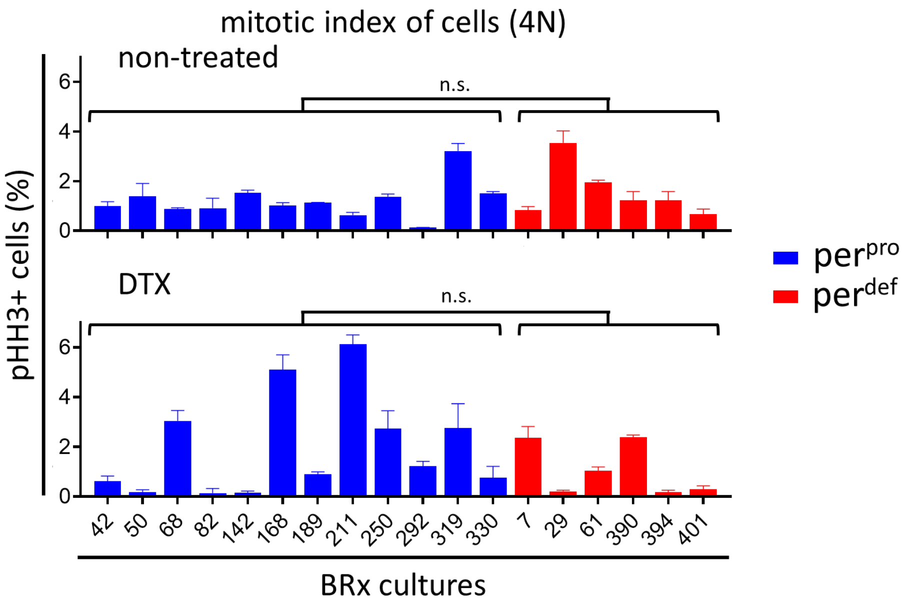

### Supplementary Figure 4

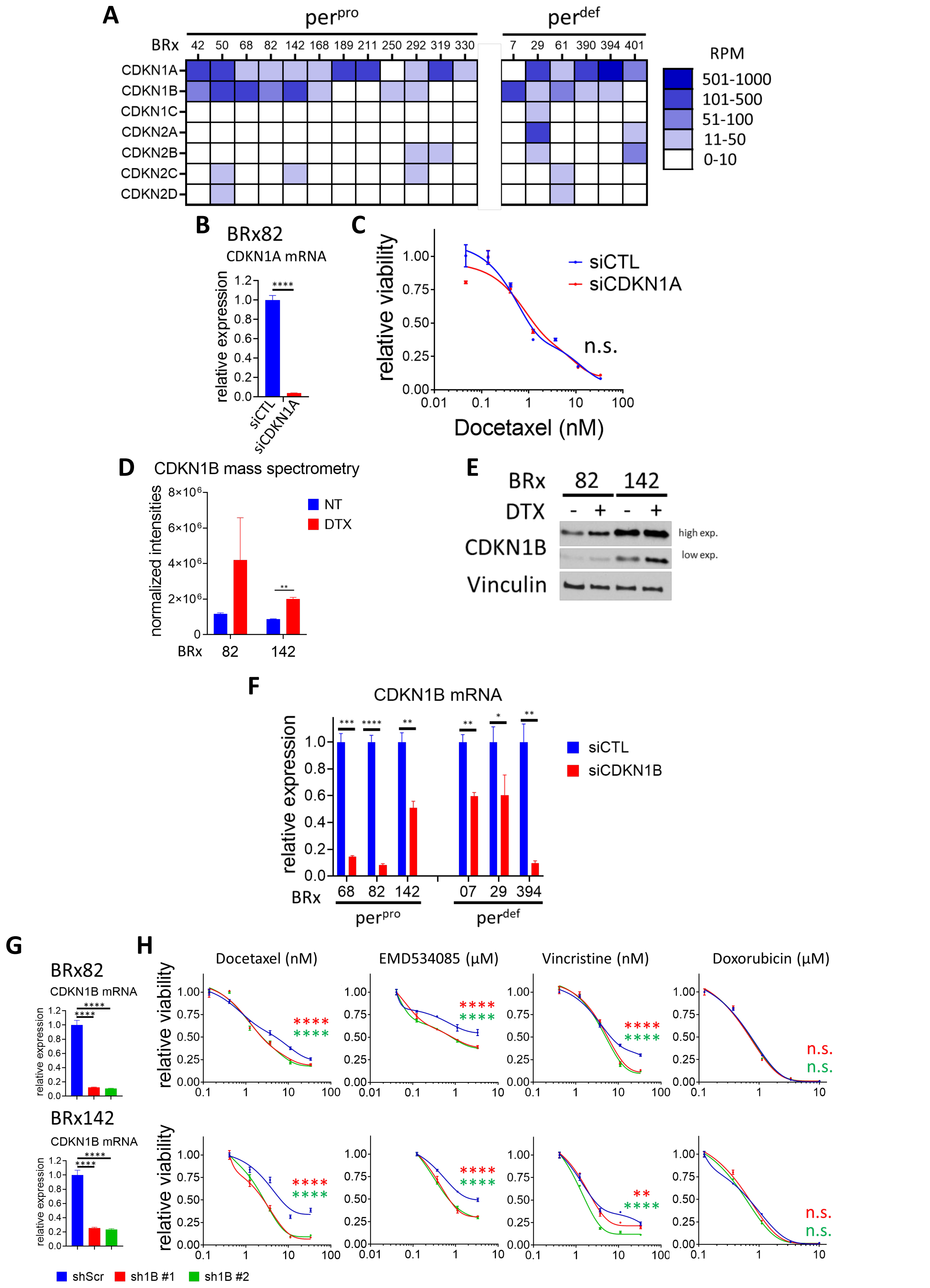

### Supplementary Figure 5

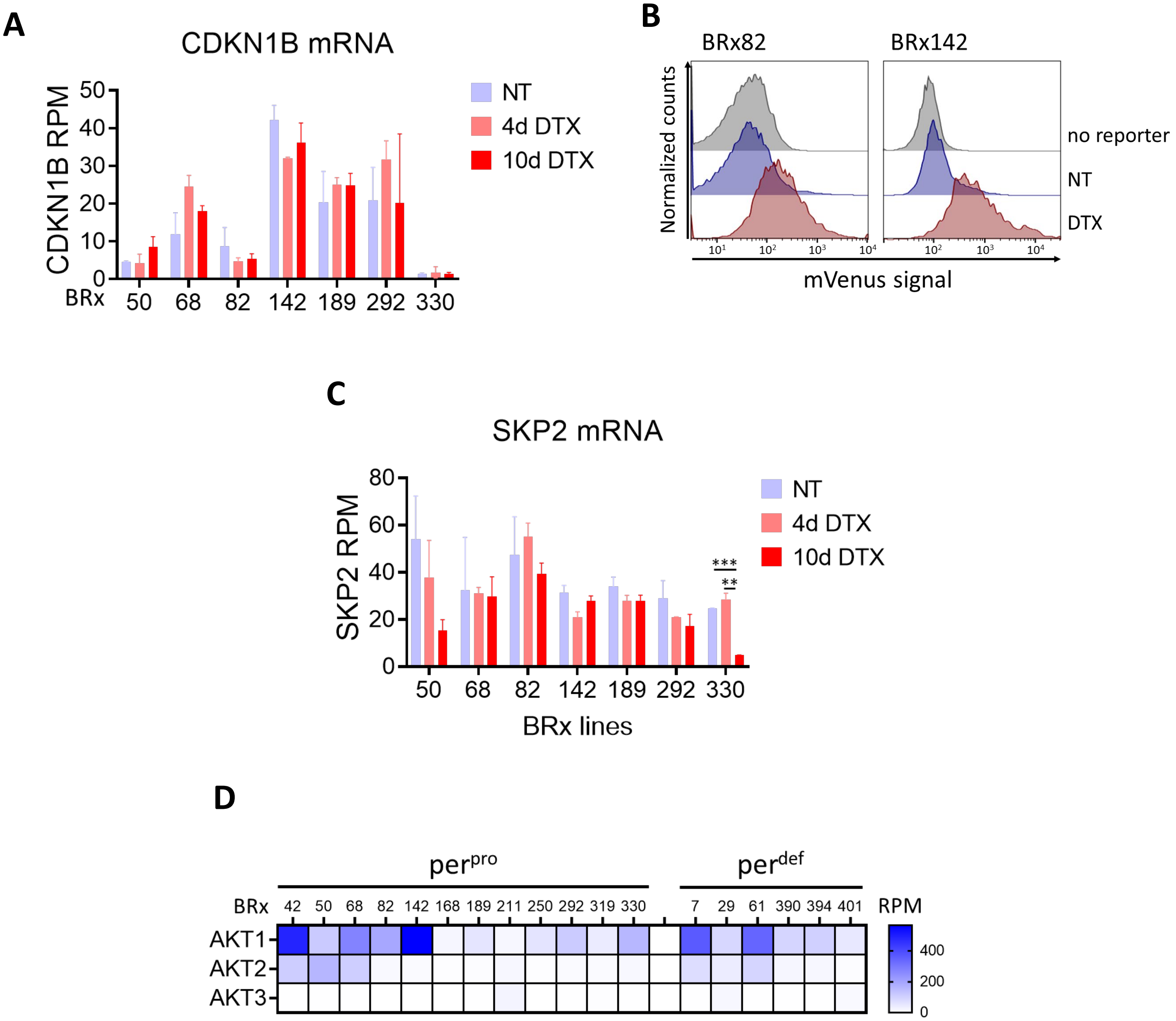
